# Single-Cell Spatial Mapping of Human Kidney Development Reveals the Critical Role of the Local Microenvironment in Cell Fate Decisions

**DOI:** 10.1101/2025.06.15.659579

**Authors:** Jonathan Levinsohn, Samuel Grindel, Bernhard Dumoulin, Amin Abedini, Aria Huang, Rena Levin-Klein, Boaz Weicz, Grace Rabinowitz, Andi M. Bergeson, Eunji Ha, Konstantin A. Klötzer, Nancy Zhang, Paul Titchenell, Mingyao Li, Joo-Seop Park, Laura S. Finn, Kotaro Sasaki, Pazit Beckerman, Oren Pleniceanu, Alex J. Hughes, Katalin Susztak

**Author notes:** **Contact Information: Katalin Susztak, MD, PhD**, Professor of Medicine, University of Pennsylvania, Perelman School of Medicine, 3400 Civic Center Blvd, Smilow Translational building 12-123, Philadelphia, PA 19104. indicates equal contribution (Jonathan Levinsohn, Samuel Grindel, Alex J. Hughes, Katalin Susztak).

## Abstract

Cell-cell interactions play a pivotal role in organ development, yet these communications have previously been studied one interaction at a time in model organisms, leaving a gap in our understanding of the cellular interplay in human development. To address this, we investigated human kidney development using single-cell RNA sequencing and spatial transcriptomics, analyzing over 500,000 cells. By mapping gene expression and differentiation trajectories in histologic space, we define the spatial organization of kidney development. Our analysis revealed newfound plasticity, showing that nephron progenitor cells undergo an early fate decision between renal corpuscle and tubular lineages. However, this choice is later reversed with some mature tubule cells transitioning back to a renal corpuscle fate. Further, through a genome-wide, spatially-aware cell-cell interaction analysis, we identified specific ligands and neighboring cell signals that create biologically meaningful cellular neighborhoods and mediate cell fate choices, offering a blueprint to understand the coordination of human development at scale.

## Introduction

Organogenesis involves intricate coordination of cell differentiation programs. This coordination is mediated by interactions between different cell types. This complex phenomenon is critical not only for proper human development, but also for understanding complex disease mechanisms and for the potential creation of artificial organs *in vitro*. Despite its significance, organogenesis remains incompletely understood. Both secreted ligands and cell-cell contact-dependent interactions play key roles in orchestrating this process.^1–4^ Cell-cell contact-dependent interactions, such as alpha-catenin signaling, are highly localized and require direct physical contact between interacting cells.^5^ In contrast, secreted ligands convey spatial information over longer distances and can form gradients.^4^ While specific examples of ligands and cell-cell contact-dependent interactions have been explored in model organisms using genetic tools, our global understanding of these mechanisms in human organogenesis is grossly incomplete.

The kidney is a spatially complex organ with over 30 specialized cell types.^6^ Its functional unit is the nephron, which comprises a filter called the glomerulus, followed by a tubule with tightly patterned segments. Human kidney development is clinically important. The number of nephrons healthy individuals are born with can vary between 200,000 and 2 million, with low nephron endowment correlating with an increased risk of developing hypertension (HTN) and chronic kidney disease (CKD).^7^ Grossly abnormal kidney development causes congenital anomalies of the kidney and urinary tract (CAKUT), which are the leading cause of CKD in children.^8^ About 15% of CAKUT cases are thought to be caused by a single gene mutation, with an additional 10% attributed to genomic copy number variations.^9,10^ Despite the clear clinical need to understand human kidney formation, research has been hampered by technical limitations.^11^

Nephron formation is highly spatially choreographed. The early steps in nephrogenesis occur in the outermost area of the developing kidney, with nephrons moving into deeper areas as they mature. Each nephron is formed from two distinct developmental lineages. The proximal part of the mature nephron—including the renal corpuscle and a large portion of the tubule—is formed from SIX Homeobox 2 (SIX2) positive nephron progenitor cells (NPCs), while the ureteric bud (UB) gives rise to the connecting tubule and collecting system.^12^ The balance between NPC renewal and differentiation is a key determinant of nephron number. Within the nephrogenic zone, NPCs form a cap of mesenchyme atop a UB tip. The earliest step of differentiation involves NPCs adjacent to the UB tip beginning to epithelialize into primitive aggregates through complex interactions with the UB. These aggregates then adopt increasingly complex, yet reproducible, spatial characteristics, forming renal vesicles, comma-shaped bodies, and S-shaped bodies.^11^ These early structures show evidence of patterning and give rise to at least five different key sub-lineages: podocytes, parietal epithelial cells (PECs), proximal tubule (PT), loop of Henle (LOH), and distal convoluted tubule (DCT). ^6,13^ Within the mature nephron, cell types follow a specific sequential spatial order—podocytes and PECs are most proximal within the renal corpuscle, followed by PT and LOH, with the DCT being the most distal. During development, the NPC- and UB-derived segments must connect to form a functional conduit for the filtered plasma.^14^ Additionally, the epithelial cells of the mature nephron must interface with other cell types from different lineages in specific locations, including endothelial cells, stromal cells, and immune cells along the tubules.

Mouse genetic studies have identified key cell-cell and ligand-receptor interactions critical for kidney development.^15^ Deletion of specific cell-cell junction proteins cause major kidney developmental defects, indicating the essential role of direct cell-cell contact.^16,17,18^ Regarding ligands, glial cell line-derived neurotrophic factor (GDNF) is secreted by the cap mesenchyme and mediates downstream cell-autonomous signaling within ureteric bud cells via the RET receptor, which is necessary for ureteric bud branching.^19^ Genetic deletion of WNT, FGF, and BMP pathway components results in defects in nephrogenesis in mice.^15^ Despite years of careful study, the full extent of these mechanisms is unclear in mouse kidney development.^15^ There is very little information on the role of ligand-receptor and direct cell-cell interactions in human kidney development. While developmental pathways are traditionally thought to be conserved in mouse models, key biological differences between the species have been noted.^13,20^

Recent studies have utilized single-cell technology to understand and describe human kidney development, shedding further light on how nephron cell types form.^21–23^ Trajectories linking progenitors to mature cell type populations are often described using “pseudotime.” Psuedotime is an ordering method that attempts to identify gene expression changes occurring during developmental transitions by taking advantage of the asynchronous development of nephrons.^24^ Studies performed in mice indicated that NPC-derived intermediates adopt renal corpuscle and tubule fates, further differentiating into podocytes and tubule cells (PT, LOH, DCT). In contrast, studies on human fetal kidney samples have proposed that intermediates take on proximal versus distal fates, with proximal cells giving rise to podocytes and PT cells.^25–27^ However, single-cell studies are unable to directly link changes in expression to histological changes within the kidney. Most importantly, the spatial context—which is critical for NPC induction and renewal—is lost during single-cell sequencing.

To address this critical issue, we integrated classic histological analysis with single-cell transcriptomics and single-cell spatial transcriptomics. We defined genetic programs associated with NPC aging and changes during differentiation and renewal. We found that there is an early decision between pre-tubular (proximal, distal, and LOH lineages) and pre-renal corpuscle (PEC, podocyte) fates. Importantly, we identified previously unreported developmental plasticity, showing evidence of maturing cell types interconverting: early PT cells becoming renal corpuscle PEC cells, and some PEC cells further transitioning into podocytes. We mapped developmental trajectories in space, identifying locations where trajectories diverge and their relationship with classic histology. Finally, we demonstrated how ligand and neighbor-mediated signaling affects human kidney development, proposed specific cell-cell interactions that cell differentiation and fate decisions. Lasty, we identified spatially distinct cellular neighborhoods defined by their surrounding microenvironment.

## Results

### Single cell RNA atlas of the developing human kidney

To understand human kidney development, we performed single cell RNA sequencing (scRNAseq) of five human fetal kidneys with gestational ages ranging from 12.5 to 20.5 weeks (**Supplementary Table 1, Supplementary Fig. 1**). After extensive data cleaning - including ambient RNA correction, doublet removal and batch correction-we generated a new atlas containing 69,170 single cells. We identified cell types from five distinct parent lineages: nephron, ureteric bud (UB), stromal, endothelial and immune cell as well as numerous differentiating, intermediate cell types (Int) (**Fig. 1a**). While some prior studies subdivided these intermediate population at this stage of the analysis, we opted against further subclustering as these cells represented a continuous gene expression and did not show clear discrete groups. We further classified nephron cell subtypes into nephron progenitor cells (NPC), podocytes, parietal epithelial cells (PECs), proximal tubule cells (PT), Loop of Henle (LOH) cells, and distal convoluted tubule cells (DCT), using classic gene markers (**Fig. 1b, c**). Cell-type specific markers (**Fig. 1b, c and Supplementary Table 2)** were consistent with the literature.^20,26–30^ Each cell type was found in every sample (**Supplementary Fig. 2 a,b**).

**Figure 1.**
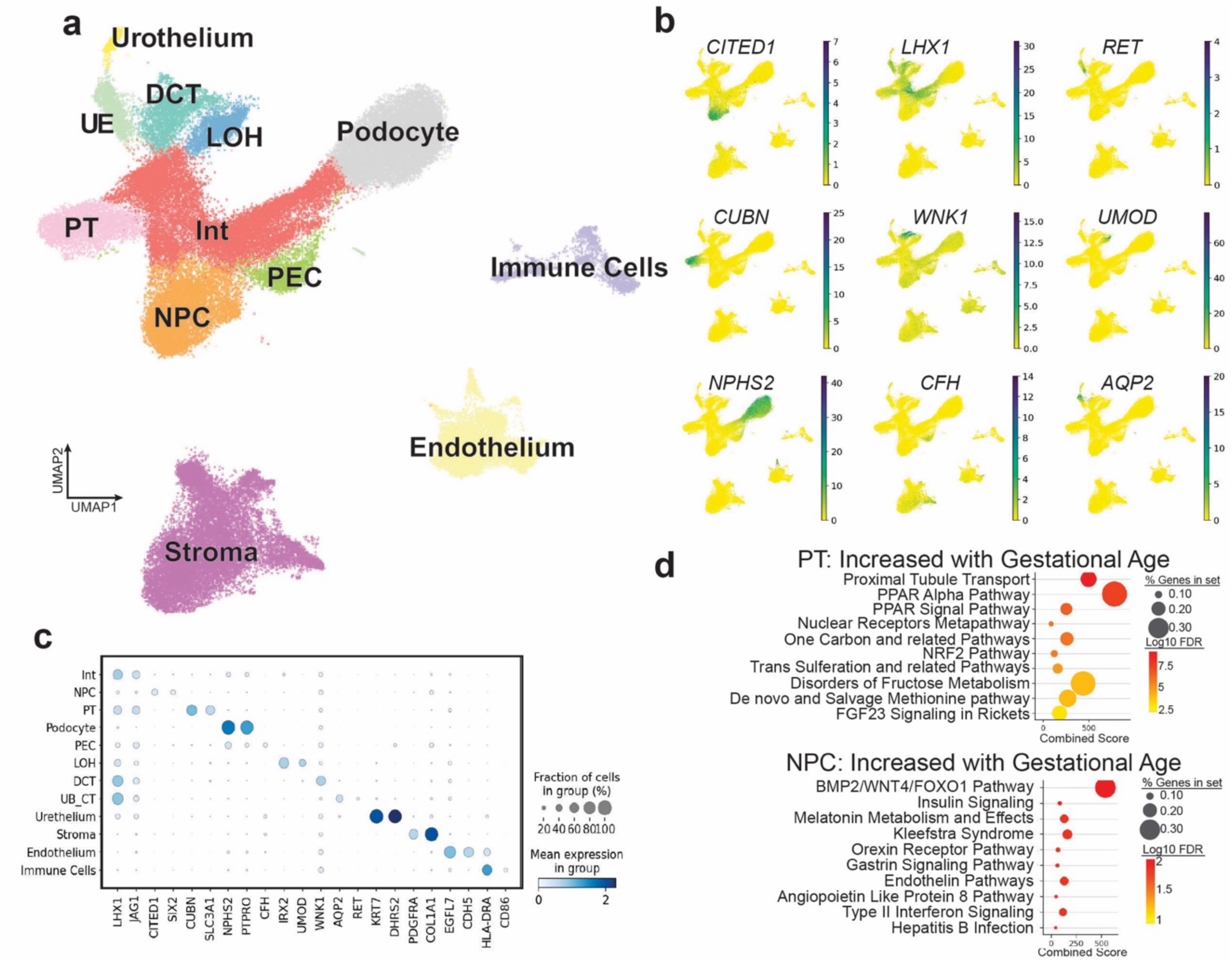
Single cell RNA atlas of the developing human kidney. (**a**) Uniform Manifold Approximation and Projection (UMAP) of single cell single cell RNA sequencing (scRNAseq) data of 65,289 cells after filtering and quality control shows populations identified within human fetal kidneys. Cell types are colored and labeled. (**b**) Feature plots showing the number of raw counts for selected marker gene (higher counts highlighted in green). The following genes are shown: Cbp/p300 interacting transactivator with Glu/Asp rich carboxy-terminal domain 1: *CITED1*, LIM homeobox 1: *LHX1*, ret proto-oncogene: *RET*, Cubilin: *CUBN*, WNK lysine deficient protein kinase 1: *WNK1*, uromodulin: *UMOD*, Podocin: *NPHS2*, Complement factor H, Aquaporin 2: *AQP2*. (**c**) Dot plot expression of representative cell type markers in each cell population. The dot size showing fraction of cells expressing the gene. The color of the dot shows mean log counts. (**d**) Pathway analysis of time dependent genes for NPC and PT cell types. Percent of genes within pathway is shown by dot size, with color indicating negative log false discovery rate (FDR). Combined score is indicated by x-axis. Int: Intermediate nephron cells; NPC: nephron progenitor cells, PT: proximal tubule, PEC: parietal epithelial cell, LOH: loop of Henle, DCT: distal convoluted tubule, UB_CT: ureteric bud/collecting duct.

NPC renewal and differentiation are key determinants of nephron number. The NPC population become exhausted during late gestation. Gestational age-associated expression changes in NPCs have functional importance in mice.^31^ We identified 467 genes with significant age dependent changes in NPCs (**Supplementary Table 3**). Pathway analysis of NPC age dependent genes showed higher Bone Morphogenic Protein 4 (BMP4)/WNT pathways, Forkhead box protein O1 (FOXO1), and insulin signaling with advancing gestational age (**Fig. 1d)**. BMP and WNT signaling are known to play a crucial role in NPC epithelialization and early nephron differentiation, indicating the correlation between gestational age and differentiation.^15^ While FOXO1 has not been shown to play a role in NPCs,^32^ FOXO1 plays an important role in insulin signaling, metabolism and also in WNT signalling.^33,34^

To understand cellular differentiation and maturation, we calculated gene expression changes associated with gestational age for each cell population (**Supplementary Fig. 2c, Supplementary Tables 9-14**). To contextualize these age-associated gene expression changes, we employed pathway enrichment analysis. Maturing PT cells showed enrichment for PT-specific solute transport, peroxisome proliferator-activated receptor (PPAR) signaling, and carbon metabolism pathways with increasing gestational age **(Fig. 1d, Supplementary Fig 2d)**. At the same time, we found that pathways associated with proliferation became less prominent with gestational age **(Supplementary Fig. 2d)**, consistent with cell type maturation.

In summary, our single cell RNAseq human fetal kidney atlas highlights key cell types and gene expression patterns. We identify increased BMP and WNT activity un aging NPCs primed for differentiation. With advancing age PT cells become metabolically active and begin expressing key cell function genes.

### Cellular differentiation trajectories of the developing human kidney

Our next aim was to reconstruct the cellular differentiation trajectories using our human fetal kidney scRNAseq dataset. RNA velocity is a method that estimates the future state of a cell based on the balance between spliced and unspliced RNA, predicting short-term future states (usually on the order of hours) by measuring the direction and speed of gene expression changes. While differentiation branchpoints are inferred from the directions of velocity vectors, RNA velocity does not explicitly identify or model them. CellRank builds on RNA velocity by adding a probabilistic framework. Instead of only predicting immediate changes, it calculates the long-term fate probabilities of cells using a Markov chain model. This approach models the entire developmental trajectory, probabilistically identifying the paths that initial cell states (e.g., progenitor cells) follow to each terminal state (e.g., differentiated cells).^35^ This is in contrast to other methods, which do not employ a probabilistic framework and determine discrete branchpoints, thereby forcing a trajectory architecture that allow for plasticity.^36,37^ This distinction is particularly important in cases, such as NPC differentiation in kidney development, where a progenitor population has multiple terminal fates.^12,22,27,28,38^

NPC progeny pattern proximal tubule and glomerular nephron segments, yet uncertainty exists over the contribution of UB-derived and NPC-derived progenitor sources for patterning distal tubule and connecting segment..^39,12^ Similar to recent studies, we observed a population of distal nephron and intermediate cells that created a continuous bridge between UB and NPC making it difficult to precisely partition these lineages (**Fig. 1a**).^20,26,27,40^ To partition these lineages in an unbiased fashion, we used CellRank to calculate the likelihood that each cell in this cluster was derived from the RET proto-oncogene (*RET)*+ UB population or the *SIX2+* NPC population. This approach partitioned intercalated cells and some connecting tubule cells to a UB lineage, while the remaining distal nephron cells were assigned to the nephron lineage (**Supplementary Fig. 1f**).^39,41^

Using CellRank, we examined cellular differentiation dynamics for the NPC lineage. We calculated the probability for each cell to transition into one of 4 final cell states (Podocyte, PT, LOH or DCT), (**Fig. 2a, b**, **Supplementary Fig. 3**), revealing clearly distinct lineages. Notably, that NPCs appear to transition to an early intermediate state and then take on either a renal corpuscle or tubular fate, consistent with observations from mouse studies (**Fig. 2b**).^27^

**Figure 2.**
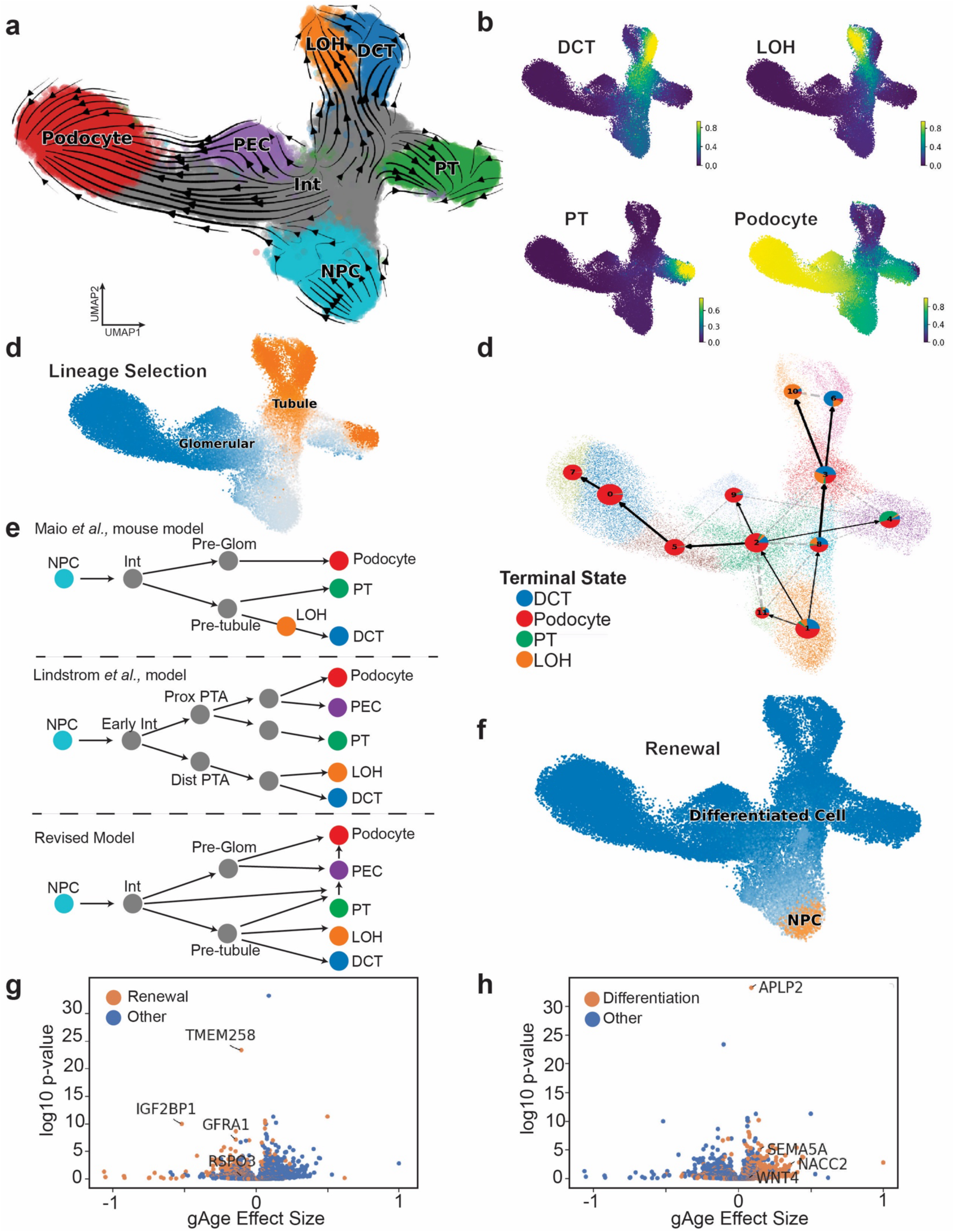
Cellular differentiation trajectories of the developing human kidney. (**a**) UMAP of nephron cell populations with RNA velocity. Colors indicate cell type annotations and arrows indicating RNA velocity direction and intensity (longer arrows indicating faster velocity). (**b**) CellRank absorption probabilities for transition from NPCs to all differentiated cell states (Podocyte, PT, LOH and DCT). Absorption probability value for each cell indicated by yellow color. (**c**) Cell rank absorption probabilities for all cells with potential terminal states of renal corpuscle (podocyte) and any tubular state (aggregate of PT, LOH or DCT). Absorption probability indicated by color on a scale of orange (high tubule probability) to blue (high renal corpuscle probability). (**d**) Directed map of NPC differentiation showing direction and connectivity of Leiden clusters. Cells are colored by Leiden clusters and each cluster has a pie chart showing an aggregate of cell fate probability of all cells within the Leiden cluster with each terminal state’s likelihood indicated by color within pie chart. Arrows indicate likely transitions with darker arrows being more frequent than thin arrows, which more frequent than dotted lines. (**e**) Schematic of differentiation from Maio et al., (Mouse development, top)^27^ Lindstrom et al.,^26^ (Human development, middle) and revised model, based on our data (Human development, bottom). (**f**) Absorption probabilities for all cells with potential terminal states of NPC and any differentiated state (aggregate of Podocyte, PT, LOH or DCT). Absorption probability indicated by color on a scale of orange (high NPC probability) to blue (high differentiated state). (**g**) Volcano plot examining gestational age (gAge) dependent expression in NPCs with renewal genes marked by orange color. Several example genes are labeled. (**h**) Volcano plot examining gestational age (gAge) dependent expression in NPCs with differentiation genes marked by orange color. Several example gene are labeled. NPC: nephron progenitor cells, INT: intermediate, PT: proximal tubule, PEC: parietal epithelial cell, LOH: loop of Henle, DCT: distal convoluted tubule. Prox PTA: proximal peritubular aggregate, Dist PTA: distal peritubular aggregate. Pre-RC: pre-renal corpuscle cell. gAge: gestational age.

To further investigate this, we calculated the probability of each cell to transition to either a tubule fate (PT, LOH or DCT) or a renal corpuscle fate (podocyte). The analysis identified populations of immature cells—referred to as intermediate cells—that had already adopted pre-renal corpuscle and pre-tubular characteristics (**Fig. 2c**). Interestingly, while many PEC cells transitioned from the pre-renal corpuscle intermediates, we found that a subset of PT cells appeared to cross over and transition into PECs (**Fig. 2c**). We observed a small number of PEC cells located near the PT cluster on the original UMAP (a few purple dots -PECs-at the bottom of the PT cluster) (**Fig. 2a, Supplemental Fig. 3**). This led us to suspect that perhaps our 2-dimensional Uniform Manifold Approximation and Projection (UMAP) did not fully represent similarities between PT and PEC. We then generated a 3-dimentional UMAP which showed the PEC population as a continuum linking PT and podocyte clusters, further suggesting that some PT may transition to PEC cells (**Supplementary Material 1**).

Next we performed a partition-based graph abstraction to further investigate these trajectories.^42^ This analysis also indicated a binary decision of NPCs becoming intermediates for the PT, LOH and DCT populations (pre-tubular), while the other cells were intermediates to podocytes and PEC (pre-renal corpuscle) (**Fig. 2d, Supplementary Fig. 3**). This NPC differentiation is fully consistent with the mouse kidney differentiation model and a recent human kidney study.^20,27^ Most interestingly, while our analysis indicated that all tubule cells shared a common progenitor, a “second path” connected PT cells and a renal corpuscle fate. This second and smaller path may explain the previously proposed sequential recruitment differentiation model (**Fig. 2e**). Again, we observed and confirmed the PT to PEC transition. To further explore, we employed a random walk analysis using CellRank. The random walk corroborated the direct conversions between PT and PECs (**Supplementary Fig. 3c**). RNA velocity data indicated that this transition is directed from PT to the PEC fate, rather than from PEC to PT (**Fig. 2a, Supplementary Fig. 3b**). Both RNA velocity, partitioned based graph abstraction, and random walk simulations observed the connections between the PECs and podocytes (**Fig. 2a**, **Fig. 2d, Supplementary Fig. 3c**). These results also indicate that the PEC cell state may be relatively unstable, with some PECs further transitioned into podocytes. Calculating fate probabilities for the UB lineage identified paths from UB progenitors to intercalated cells, urothelium and connecting tubule (**Supplementary Fig. 4**).

We then sought to identify genes that drive cellular differentiation for each trajectory. CellRank performs this by correlating absorption probability for a given trajectory with each gene’s expression. We performed this calculation for tubular and renal corpuscle cell fates (**Supplementary Table 10**) and for each terminal state (DCT, PT, LOH and podocytes) individually (**Supplementary Table 11**). Amongst other genes, we identified Iroquois homeobox 1 (*IRX1*) for the LOH trajectory. *IRX1* is thought to be important to be important driver nephron morphogenesis.^43^ However, we also noted that many of these genes were markers of the mature cell type and perhaps less likely to be actively mediating cell fate, such as podocin (*NPHS2*) for podocytes.

NPC renewal and differentiation is critical for nephron formation. To better understand factors that govern NPC renewal using CellRank to calculate absorption probabilities estimating the likelihood of a cell becoming an NPC compared to the likelihood of that cell becoming any differentiated state (PT, podocyte, LOH or DCT) (**Fig. 2f**). We identified genes correlated with each of these absorption probabilities to pinpoint genes associated with renewal (**Supplementary Table 12**). Supporting the validity of these renewal-associated genes (**Supplementary Fig. 2a**), we found that genes associated with renewal also tended to have higher expression in earlier gestational ages (**Fig. 2g, Supplementary Fig. 5**). Conversely, expression of differentiation associated genes were higher at later gestational ages (**Fig. 2h, Supplementary Fig. 5**).

In summary, using a probabilistic cell differentiation model and partition-based graph abstraction, we found strong evidence that NPC-derived intermediates take on both pre-tubule and pre-renal corpuscle fates prior to terminal differentiating. Importantly, we detect previously unrecognized plasticity in PT intermediates, which can transition into PEC cells and subsequently into podocytes.

### Spatial molecular characterization of the developing human kidney

Cells are presumed to interpret their position within a tissue to make critical decisions about their fate, ensuring that complex tissues and organs are organized correctly and function optimally. This spatial context awareness of cells has been poorly understood. An excellent example of this is the proximity of UB to NPC cells, drives cellular differentiation. To understand how cellular locations impact differentiation in human fetal kidneys, we performed spatial transcriptomics on 3 human fetal kidneys with gestational ages of 15 to 19 weeks using the CosMx platform,^44^ and characterized the expression of 1,000 genes at single-cell resolution (**Supplementary Fig. 7a, Supplementary Table 1**). We filtered and performed rigorous quality control (QC) to characterize over 475,000 cells in our spatial atlas (**Supplementary Fig. 6a, Supplementary Table 1**). We generated a UMAP of the CosMx reads, clustered cells, and annotated cell types. All spatially detected cell types were also present in our dissociated single-cell data (**Supplementary Fig. 6b).** We next used variational autoencoders, scVI and scANVI,^45^ to integrate the scRNA-seq/CosMx datasets (**Supplementary Fig. 7a, b**). This combined integration enabled the imputation of whole transcriptome gene expression for spatial data. We did this by mapping each CosMx cell to the 15 most similar scRNAseq cells and calculated a weighted average for each gene (**Supplementary Fig. 7c-e).** Furthermore, we performed this mapping and imputation for CellRank absorption probabilities, which allowed us to examine cell trajectories in space (**Supplementary Fig. 7e)**.

In this integrated atlas, we found discrete cell types on a shared UMAP (**Fig. 3a**) with annotations being conserved from the original (pre-integrated) CosMx annotations (**Supplementary Fig. 8 a,b**). Furthermore, spatial locations of cell types were consistent with known tissue organization and histology. For example, tubule cell types mapped to tubule morphologies, and morphologic renal corpuscles contained podocytes, endothelial cells and PECs (**Fig. 3b, Supplementary Fig. 9**). Using the integrated dataset, we plotted the spatial locations of gene expression for all genes in the developing human kidney samples (**Fig. 3c, d, Supplementary Fig.10**). We found early NPC markers such as *UNC Homeobox (UNCX)* localized to the outermost portion of the tissue. With increasing tissue depth, we found higher expression of marker genes for intermediate cell states such as *LIM Homeobox 1* (*LHX1)* and *Jagged canonical notch ligand 1 (JAG1)* (**Fig. 3d**). Deeper still, we identified markers of trajectory endpoints, such as apolipoprotein E (*APOE),* which marks mature PT cells (**Fig. 3d**), and *podocin (NPHS2),* which marks mature podocytes (**Fig. 3c**).

**Figure 3.**
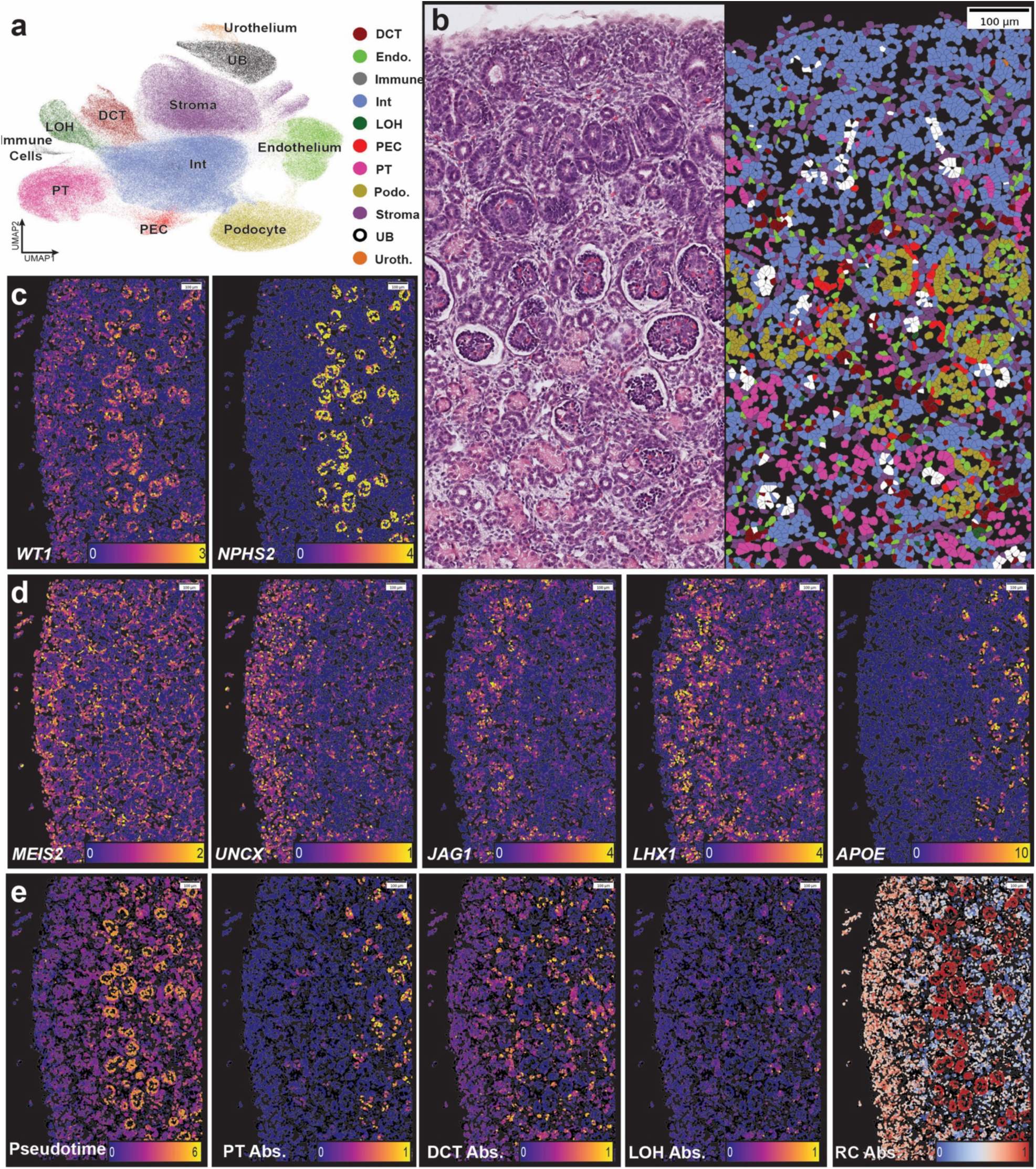
Spatial molecular characterization of the developing human kidney. (**a**) Integrated UMAP of spatial (CosMx) and dissociated scRNAseq data showing cell type annotations indicated by color. (**b**) H&E-stained human fetal kidney section adjacent to the corresponding cell type annotation from our spatial transcriptomic data. Cell type annotation is indicated by color. (**c**) Expression of Wilms tumor antigen 1: *WT1*, podocin: *NPHS2* (podocyte marker genes in yellow) in our analyzed kidney sample. (**d**) Expression of Meis homeobox 2: *MEIS2*, UNC homeobox: *UNCX*, LIM homeobox 1: *LHX1*, Apolipoprotein E: *APOE*, Jagged canonical Notch ligand 1: *JAG1*, (yellow) in the developing human kidney. (**e**) Pseudotime of NPC lineages, colored from early (purple) to late (yellow); absorption probabilities for PT, DCT and LOH colored from low (purple) to high (yellow); Renal corpuscle fate absorption probabilities colored from low (blue) to high (red), within the developing human kidney. PT: proximal tubule, DCT: distal convolutes tubule, LOH: loop of Henle, RC: renal corpuscle cells.

Next, we aimed to visualize the kidney developmental trajectories in space. Previously development was only ascertained by observing certain structures (comma- or S-shaped bodies); however, these structures may not represent important developmental milestones. The fully integrated gene expression dataset enabled the visualization of developmental and differentiation stages—as absorption probabilities—in our histological images (**Fig. 3e**). This analysis confirmed the known cortic-medullary spatial organization associated with kidney development; the most developmentally immature nephron cells (lowest values for pseudotime) were located in the most superficial portion of the tissue, while more differentiated stages were located deeper.^13^ More interestingly, the quantitative nature of our analysis permitted us to identify where individual cell fates appear to diverge. We found podocyte and DCT specification in a relatively superficial region, indicating an early lineage specification consistent with our CellRank analysis (**Fig. 3e**). Similarly, plotting the renal corpuscle absorption probability onto the spatial gene expression data, indicated an early commitment of intermediate cells onto the renal corpuscle fate (**Fig. 3e**).

In summary, by integrating whole-transcriptome scRNAseq and single-cell spatial transcriptomic data, we generated a single cell resolution map of the developing human kidney. Integration enabled the mapping of developmental stages and trajectories onto histological slides, generating a new appreciation for kidney development.

### Molecular characterization of the histologic neighborhoods in the developing human kidney

Human fetal kidney development has been traditionally described by specific structures such as the blastema (the region containing the most undifferentiated cells), primitive epithelial structures (PES), which consist of the renal vesicle, comma and S-shaped bodies, and early renal corpuscles. Previous scRNA-seq analysis performed in dissociated kidneys had difficulties matching gene expression changes to these histological and physiological hallmarks, as intermediate cells present as a continuous cell population rather than discrete cell types. With the combination of spatial transcriptomics and single cell sequencing, we aimed to understand how gene expression and differentiation trajectories relate to these classic anatomic structures. We had an expert pathologist annotate these developmental structures (**Fig. 4a, b, Supplementary Fig. 11a)**, and we examined gene expressions, cell type fractions and trajectories for each neighborhood using our spatial gene expression information.

**Figure 4.**
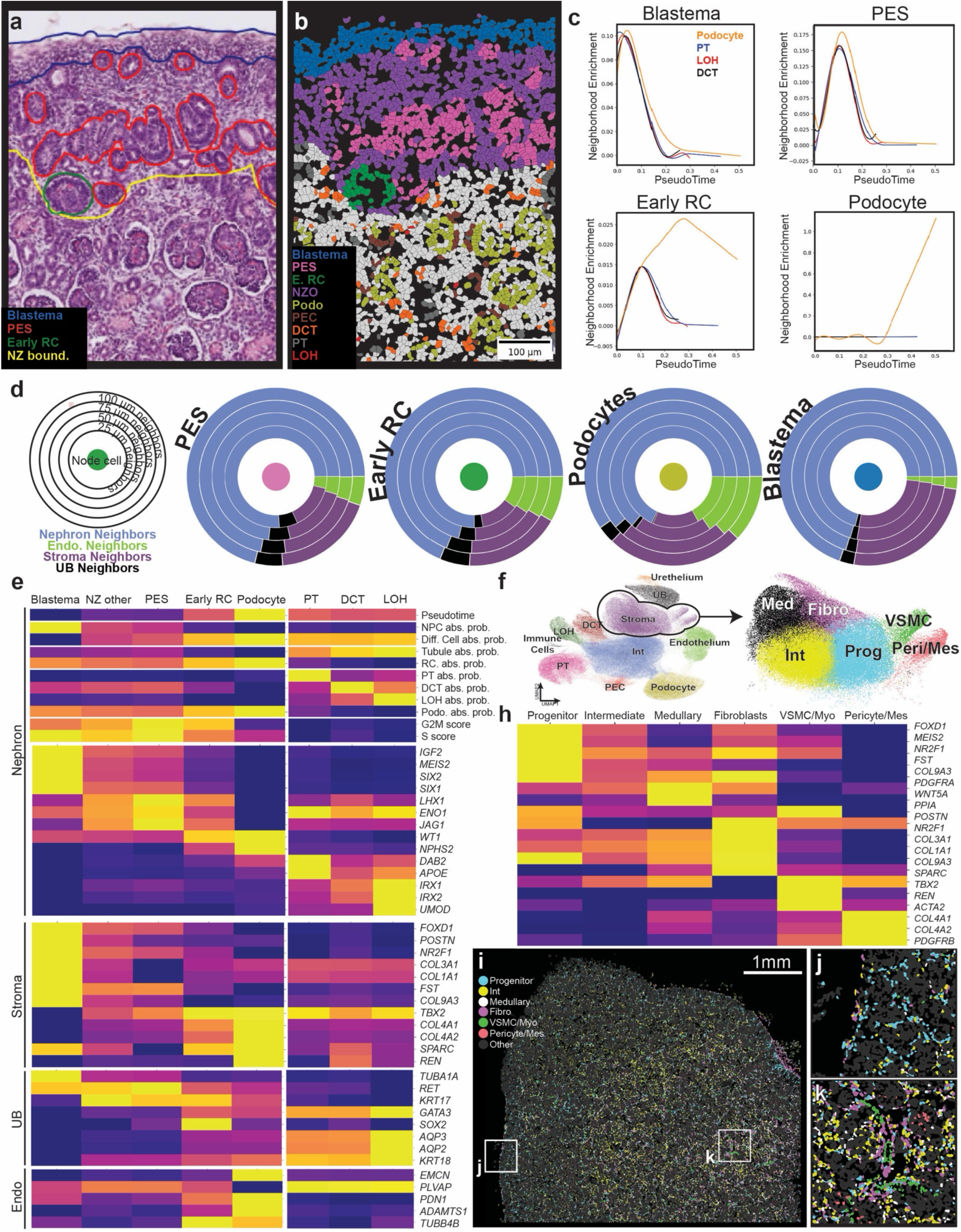
Molecular characterization of the histologic neighborhoods in the developing human kidney. (**a**) H&E image of the nephrogenic zone in a human fetal kidney with manual neighborhood annotation. Blastema is circled by blue lines, primitive epithelial structures (PES) in red, and green shows early renal corpuslces (E. RC), with the lower boundary of the nephrogenic zone outlined in yellow. (**b**) The H&E identified neighborhoods were then plotted on the CosMx data as shown in a small section and across the entire tissue, with cells colored by neighborhood. (**c**) Pseudotime trajectory characteristics of each neighborhood. For each histologic neighborhood, we graph aggregate trajectory enrichment (y-axis) against pseudotime (x-axis) for each linage, as indicated by color. Remaining neighborhoods and cell types are presented in Supplementary Fig. 10. (**d**) Neighbor characteristics of each histologic neighborhood. Circular plots show node cell, and neighbor frequency with increasing neighbor radius. Neighbor type is indicated by color. Remaining neighborhoods and cell types are presented in Supplementary Fig. 10. (**e**) Neighbor frequency across pseudotime. For each neighbor type (stroma, UB and endothelium), we plot neighbor number (y-axis) against pseudotime, for each trajectory, which are colored individually. (**f**) Aggregate trajectory and expression characteristics of each histologic neighborhood across lineages. Matrix plot showing gene expression or differentiation characteristics within each neighborhood as parsed by lineage. Each column shows a histologic neighborhood and each row a specific gene or trajectory. Value is indicated by color with low being purple and high being yellow. (**g**) UMAP of integrated single cell RNAseq and CosMx, showing cell annotations. Stromal cells (circled) were subclustered and these subclusters were annotated as shown on adjacent UMAP of just stromal cells. Subclusters are marked by color. (**h**) Aggregate expression characteristics of each stromal subcluster. Matrix plot showing gene within each neighborhood as parsed by lineage. Each column shows a subcluster and each row a specific gene. Value is indicated by color with low being purple and high being yellow. (**i**) Stromal subclusters in space. Stromal cells are plotted in space with annotation of each cell type indicated by color. Insets J and K are marked by a labeled white box. Grey cells indicate non-stromal cells. (**j**) Stromal subclusters in space, inset of panel I, showing a region of nephrogenic zone. Stromal cells are plotted in space with annotation of each cell type indicated by color. (**k**) Stromal subclusters in space, inset of panel I, showing a deeper region within the kidney. Stromal cells are plotted in space with annotation of each cell type indicated by color. PT: proximal tubule, LOH: loop of Henle, DCT: distal convoluted tubule, PEC: parietal epithelial cell, Early RC: early renal corpuslce, PES: primitive epithelial structure, NZO: nephrogenic zone other, UB: ureteric bud.

We first examined how trajectories correlated with these histologic. Blastema showed the earliest pseudotime and no enrichment for a particular cell fate, consistent with the notion that this neighborhood houses stem cells (**Fig. 4c**). PES demonstrated a later pseudotime, but still no enrichment for a particular cell fate, showing that the next developmental stage can still give rise to all downstream lineages (**Fig. 4c**). In contrast, early renal corpuscles, had an even later pseudotime and an enrichment for the podocyte lineage, demonstrating that a fate decision has been made within this neighborhood (**Fig. 4c**). Our analysis was corroborated by mature podocytes demonstrating full enrichment for the podocyte lineage and the latest pseudotime.

Recognizing that these fate decisions are likely mediated by neighboring cells, we quantified the cell types present in each histologic neighborhood (**Fig. 4d**). To achieve this, we analyzed the registered cells within 25-100 uM of the center cell. Cells within the blastema were almost exclusively surrounded by other NPC and stromal cells, UB and endothelial cells were exceedingly rare in the blastema. This region likely represents the self-renewing NPC cells. PES had the highest fraction of UB cells, lower stromal and higher endothelial cells. This is consistent with observations that NPC differentiation occurs at this stage in close proximity to the ureteric bud. Endothelial cell fraction further increased in early renal corpuscles and podocytes, suggesting that endothelial cells may be important in establishing podocyte fate.

We next examined cell-type gene expressions within each neighborhood. **Figure 4e** shows the expression of genes within the NPC, UB, stromal and endothelial cells within each developmental structure: blastema, PES, early renal corpuscle, and different tubule segments. The gene expression data indicated remarkable gene expression variability for the same cell type within different structures (**Fig. 4e**). For example, the early interemediate cells still expressed SIX1/2 in the blastema, but gradually express markers of differentiated cells, podocytes, PT, LOH etc. in different neighborhood contexts. Similarly, UB expressed RET in the blastema and AQP2/3 in differentiated PC cells. Interesting to note that while endothelial cells started off with a common progenitor, they showed important differences in the glomerulus (expressing *EDH3*) or peritubular endothelial cells (expressing *PLVAP*). While it is recognized that there is stromal heterogeneity in both the adult and fetal kidney, linking expression to spatial location has not been examined beyond examination of single marker genes.^30,46^ Stromal cells were very common in the blastema and these cells were strongly positive for FOXD1 (**Fig. 4f,g, Supplementary Fig. 12**). They formed a nest around the differentiating NPCs. We then further subclustered stromal cells, which revealed biologically and spatially distinct stromal populations. We identified progenitor cells, intermediate cells, medullary fibroblasts, pericytes, mesangial cells, fibroblasts vascular smooth muscle cells and myofibroblasts, and examined marker gene expression as well as location (**Fig. 4g-j, Supplementary Fig. 12**). FOXD1-positive progenitors were located mostly in the nephrogenic zone, WNT5A-positive medullary stroma was within the medulla, and vascular smooth muscle/myofibroblasts appeared in vessel-like shapes, and pericytes/mesangial cells were located within renal corpuscles (**Fig. 4g-j, Supplementary Fig. 12**). These results indicate the key role of spatial location in explaining gene expression variation, much of which correlates with histologic architecture.

In summary, we found that our integrated single cell spatial gene expression and cell trajectory analysis strongly correlated with classic developmental structures observed on histological slides. We provide a gene expression, cell interaction and developmental trajectory map for these classic developmental programs observed in human kidneys, including a detailed analysis for stromal cells.

### Spatially-resolved, unbiased cell-cell interactions in the developing human kidney

Cells have a unique ability in the body to obtain spatial information. Ligand-receptor interactions play a key role in enabling cells to interpret spatial cues. Using our ability to define cell differentiation trajectories in space, we aimed to identify specific cell-cell interactions that regulate these trajectories in an unbiased manner. Traditional ligand-receptor interaction analyses are limited in scRNA-seq analysis by the lack of spatial proximity information. Our new spatial gene expression data enabled a spatially aware ligand-receptor analysis, with full-transcriptome estimated expression permitting widespread discovery.

First, we used CellPhoneDB^47^, restricting interactions to those found within histologically identified neighborhoods (**Supplementary Table 13**). CellPhoneDB highlighted 569 spatially plausible interactions, including classic ligand-receptor interactions such as WNT (Wingless/Integrated), FGF (Fibroblast Growth Factor), and BMP (Bone Morphogenic Protein) ligands, which are known to be critical for kidney development in mouse genetic experiments (**Supplementary Table 13**). While CellPhoneDB was effective, it requires predefined neighborhoods, which may or may not correctly discriminate cells that can plausibly interact from those that cannot.

We next employed CytoSignal^48^ which leverages spatially resolved gene expression data to calculate signaling among single cells at precise spatial coordinates, not limited to previously defined structures. CytoSignal accounts for ligand diffusion in identifying ligand-receptor interactions and also identifies direct cell-cell interactions by requiring signaling cells to be in direct contact. In the developing human kidney, we identified approximately 300 interactions per sample, with 246 of these interactions conserved across all three samples (**Supplementary Table 14**). CytoSignal revealed key signaling nodes such as Wnt family member 4 (WNT4), which coordinates early differentiation resulting in epithelialization of nephron progenitors in mice.

Interestingly, both CellPhoneDB and CytoSignal indicated blastema-specific insulin-like growth factor 2 (IGF2) signaling, with the ligand generated in both nephron progenitor cells (NPCs) and stromal cells, and the insulin-like growth factor 1 receptor (IGF1R) expressed throughout the kidney (**Supplementary Fig. 13**). While CytoSignal highlighted the spatial location of ligand-receptor interactions, it could not indicate the biological impact of these interactions. To address this, we examined individual sender signals—combinations of ligands and direct cell-cell neighbor-dependent signals—for each cell (**Fig. 5a**). We then calculated correlations between sender signals (e.g. ligands) in the CytoSignal database and each trajectory (**Fig. 5b**). This analysis allowed us to identify cell-cell interactions associated with specific biological processes. We found that Glial Cell Line-Derived Neurotrophic Factor (GDNF) and R-spondin 3 (RSPO3) showed a high correlation with nephron progenitor cell (NPC) renewal (**Fig. 5b**). Furthermore, the presence of IGF2 appeared to be associated with an early stem cell state across all NPC lineages (**Fig. 5c**). We found that the cognate receptors for IGF2 were constitutively expressed in NPC lineage cells (**Fig. 5d, Supplementary Fig. 13**), while the ligand was produced by NPC cells, as well as stromal and ureteric bud (UB) cells. This led us to hypothesize that downstream IGF2 signaling might have a specific non-cell-autonomous role in NPC function.

**Figure 5.**
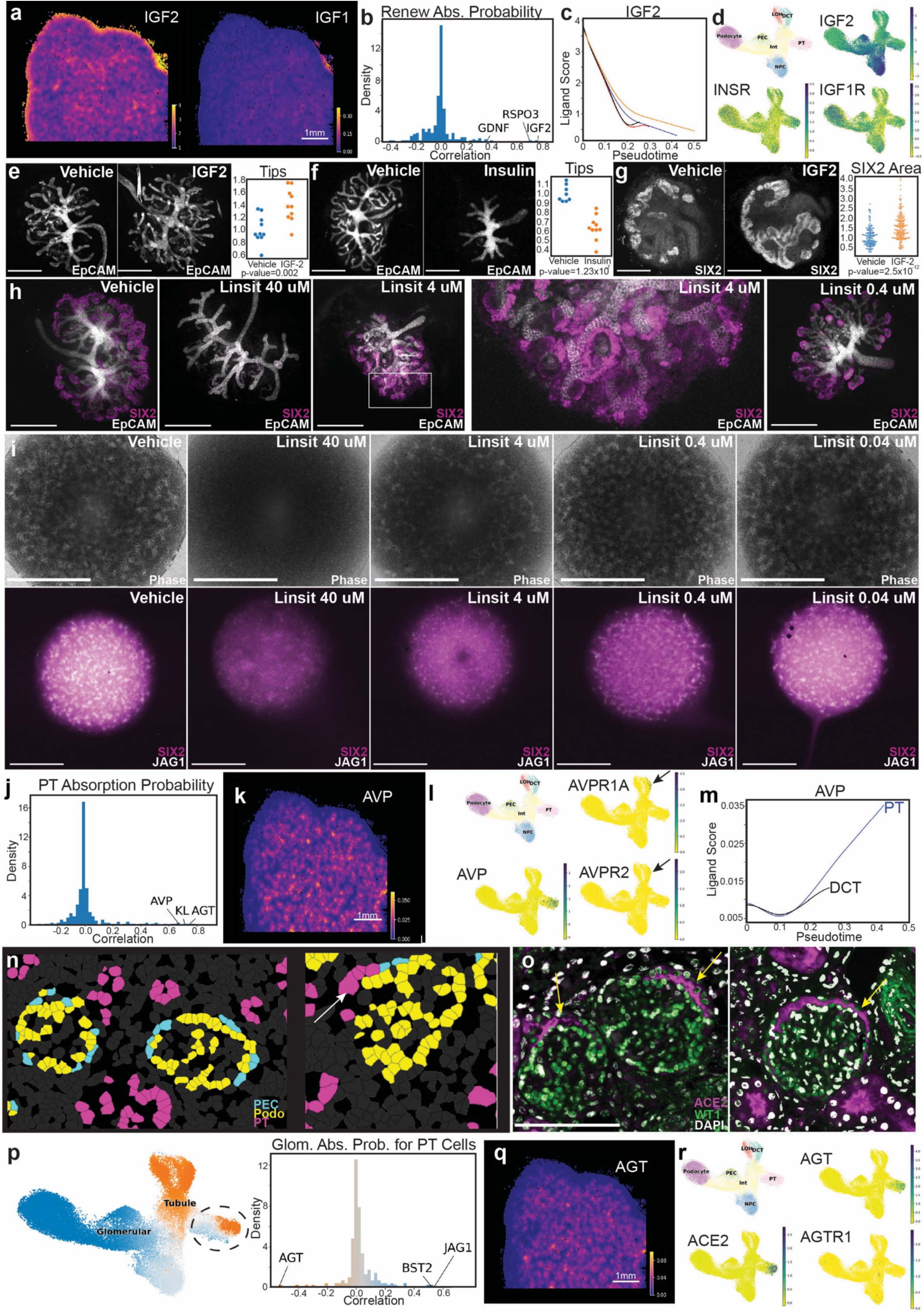
Spatially-resolved, unbiased cell-cell interactions in the developing human kidney. (**a**) Spatial locations of CytoSignal ligand scores for IGF1 and IGF2 in the human kidney, with score indicated by color (low being purple and high being yellow). (**b**) Histogram of correlations of microenvironment score with absorption probability for 878 microenvironment signals for the renewal trajectory. X-axis shows correlations and y-axis shows frequency (density estimate) of that correlation. Some individual genes of interest are labeled. (**c**) Ligand score (y-axis) is plotted against pseudotime (x-axis) for IGF2. Each trajectory is shown by its discrete color. (**d**) UMAP of scRNAseq showing That IGF2 is most expressed in the NPC cell population, with cognate receptors IGF1R and INSR expressed in all cell populations. (**e**) EpCam immunofluorescence staining (white: ureteric tree) images and quantified number of UB tips (y-axis) for mouse fetal explants incubated with ectopic IGF2 ligand. Total number of tips was normalized for average number of tips in the control samples for each replicate. This was performed in 3 separate litters. Scale bar = 400 um.} (**f**) EpCam immunofluorescence staining (white: ureteric tree) images and quantified number of UB tips (y-axis) for mouse fetal explants incubated with ectopic insulin ligand. Scale bar = 400 um. This experiment was repeated in a separate mouse litter. Total number of tips was normalized for average number of tips in the control samples for each replicate. This was performed in 3 separate litters. Scale bar = 400 um. Each replicate shown separately in Supplementary Fig. 15. (**g**) SIX2 immunofluorescence staining (white: NPC marker) images and quantified size of SIX2 caps. Area was normalized to average area for control explants for each 3 kidneys per condition performed in 2 separate litters (12 total kidneys). Scale bar = 400 um. Each replicate shown separately in Supplementary Fig. 15. (**h**) SIX2 (purple) and EpCam (white) immunofluorescence of explants treated with IGF1R inhibitor linsitinib at various concentrations. This was repeated in a separate litter and had similar results. Scale bar = 400 um. Each replicate shown separately in Supplementary Fig. 15. (**i**) Phase, SIX2 and JAG1 immunofluorescence of human kidney organoids treated with IGF1R inhibitor linsitinib at various concentrations. This experiment was repeated with similar results in a separate organoid batch. Scale bar indicates 1000 um. (**i**) Histogram of correlations of microenvironment score with absorption probability for 878 microenvironment signals for the PT trajectory. X-axis shows correlations and y-axis shows frequency (density estimate) of that correlation. Some individual genes of interest are labeled. (**j**) Spatial locations of CytoSignal ligand scores for AVP in the human kidney, with score indicated by color (low being purple and high being yellow). (**k**) UMAP of scRNAseq showing That AVP is most expressed in the PT cell population, with cognate receptors AVPR1A and AVP2R expressed in the DCT cell population. (**l**) Ligand score (y-axis) is plotted against pseudotime (x-axis) for AVP. DCT and PT trajectory are shown by their own discrete color. (**o**) CosMx cell type annotations of renal corpuscles, with PT, podocyte and PEC annotations shown in space. (**p**) Immunofluorescence of human fetal kidney tissue showing ACE2 expression (purple), WT1 expression (green) and DAPI (white) (**q**) UMAP showing probability of trajectories for renal corpuscle and tubular fate indicated for each cell based on color (left). Histogram of ligand correlations with renal corpuscle fate within the PT population. X-axis shows correlations and y-axis shows frequency (density estimate) of that correlation. Individual genes of interest are labeled. Scale bar = 100 uM. (**r**) Spatial locations of CytoSignal ligand scores for AVP in the human kidney, with score indicated by color (low being purple and high being yellow). (**s**) UMAP of scRNAseq showing That AGT, and ACE2 are expressed in the PT cell population, with cognate receptor AGTR2 expressed in all NPC derived cell types. PT: proximal tubule, LOH: loop of Henle, DCT: distal convoluted tubule, PEC: parietal epithelial cell, NPC: Nephron progenitor cell, Int: intermediate. IGF2: insulin like growth factor 2, AVP: arginine vasopressin, IGF1: insulin like growth factor 2. AVP: Arginine vasopressin. AGT: Angiotensinogen, ACE2: Angiotensin converting enzyme 2, AGT2R: Angiotensin 2 receptor, AVPR1A: arginine vasopressin receptor 1a, AVPR2: arginine vasopressin receptor 2. MsE: mouse explant. HumSpatial: Human Spatial, HumOrg: Human organoids, HumIF: human immunofluorescence.

To investigate this, we employed a mouse fetal explant system and cultured kidneys from embryonic day 13 (E13) embryos. Mouse fetal kidney explants were exposed to ectopic mouse IGF2, which resulted in more ureteric bud (UB) tips after five days of *ex vivo* culture compared to controls, indicating that IGF2 increases UB branching (**Fig. 5e, Supplementary Fig. 15a**). IGF2 is part of the insulin signaling pathway and canonically acts on the IGF1 receptor (IGF1R) but can also bind to the insulin receptor. Insulin can also bind IGF1R, especially at higher concentrations, as observed in clinical situations like maternal diabetes. However, cultured explants treated with high-dose insulin showed poor growth and decreased UB branching relative to controls (**Fig. 5f, Supplementary Fig. 14b**). Our immunohistochemistry analysis also indicated positive phospho-AKT staining—a canonical marker of insulin signaling—in the blastema and nephrogenic zone (**Supplementary Fig. 15**). To specifically test the effect on NPCs, we performed immunofluorescence on IGF2-incubated explants and found that caps were larger than in the control group (**Fig. 5g, Supplementary Fig. 14c**). Furthermore, incubation with an IGF1R inhibitor, Linsitinib, led to the loss of a SIX2+ population at high concentrations and smaller, disorganized caps at lower concentrations (**Fig. 5h**). We then examined NPC lineage human kidney organoids using a previously generated snRNAseq dataset capturing multiple time points.^49^ These data showed that IGF1R, IGF2R, and INSR receptors were highly expressed, with early IGF2 expression present at day 12, but decreasing by day 26 (**Supplementary Fig. 16**). This prompted us to examine whether insulin signaling was critical for human organoid development. Indeed, treatment with linsitinib showed decreased SIX2-positive cells and fewer tubular aggregates at day 12 with increasing linsitinib concentrations (**Fig. 5i**). Conversely, adding IGF2 modestly increased both SIX2- and JAG1-positive populations at 12 days (**Supplementary Fig. 17**).

We found that arginine vasopressin (AVP) and angiotensinogen (AGT) had a high correlation with the proximal tubule (PT) lineage (**Fig. 5j**), with more mature PT cells expressing the ligand at higher levels. AVP ligand was observed to be localized to mature tubular areas within the tissue (**Fig. 5k**), and the AVP gene was exclusively expressed by PT cells (**Fig. 5l**). However, cognate receptors for AVP were not found in PT cells (**Fig. 5l**), but were present within the distal convoluted tubule, suggesting that AVP does not act upon the cells that secrete it. Classically, AVP is produced by the hypothalamus; however, recent studies have shown that some local AVP production also occurs in the adult kidney, though its function is somewhat unclear.^50^ Our spatial data shows that AVP is spatially associated with the PT trajectory, which produces the ligand, and the DCT trajectory which expresses the cognate receptors (**Fig. 5m**). This suggests that local AVP production in the kidney may act to temporally synchronize differentiation across nephron segments. This is supported by previous work on the ureteric lineage which showed that organoids had improved differentiation of intercalated cells (IC) and principal cells (PC) with the addition of ectopic AVP.^51^ To further examine this in the NPC lineage, we used human iPSC-derived organoids. After finding that AVP and its receptors were present in NPC-derived organoids (**Supplementary Fig. 16**), we hypothesized that AVP had a similar pro-differentiation effect on NPC-derived DCT cells. We found that an inhibitor of AVP receptors (tolvaptan) led to fewer and smaller DCT segments, while AVP led to larger DCT segments (**Supplementary Fig. 18 a, b**).

With an improved understanding of cellular spatial context, we re-examined the newly predicted PT to parietal epithelial cell (PEC) transition. We examined our spatial transcriptomics data to see if these predicted transitions could be visualized, finding that most renal corpuscles were composed of PECs and podocytes, in addition to stromal and endothelial cells. However, in a small fraction of renal corpuscles, we noted that the corpuscle appeared to be lined with PT cells (**Fig. 5n**). To further examine this, we performed immunofluorescence on human fetal kidney tissue and found that some renal corpuscles were lined with ACE2-positive cells (a marker of PT cell identity), suggesting that these morphologic PECs had molecular characteristics of PT cells, supporting the predicted transition (**Fig. 5o, Supplementary Fig. 19**). Examining adult human tissue and postnatal day 0 (P0) mouse tissue, we did not identify such a pattern, suggesting that this may be specific to the human fetal state (**Supplementary Fig. 19**). To examine how this plasticity might be controlled, we investigated which ligands were associated with the PT to PEC transition (**Fig. 5q**), identifying that Angiotensinogen (AGT) was strongly associated with PT cells maintaining their PT identity, while JAGGED1 (JAG1) and BST2 were associated with transition to PEC. Given the high levels of renin and ACE2 in the developing kidney, we suspected that angiotensin II, the product of angiotensinogen, might be important in this transition. Alterations in ACE2 and AGT are known to cause abnormal kidney tubule development in both mice and humans; however, the mechanism is unclear.^52,53^ We treated organoids with AGTII, finding increased PT lineage cells and decreased podocyte lineage, further supporting the notion that AGT may act as a feed-forward mechanism establishing PT fate (**Supplementary Fig. 18c, d**).

We performed ligand association analysis for the other NPC trajectories, finding that Fibroblast Growth Factor 9 (FGF9), Kininogen 1 (KNG1)—a precursor to bradykinin—and cadherin-1 (CHD1) were correlated with the loop of Henle (LOH) trajectory, while CHD1, Ephrin A4 (EFNA4), and Bone morphogenic protein-3 (BMP3) were associated with the distal convoluted tubule (DCT) (Supplementary Fig. 20). We also identified Vascular Endothelial Growth Factor A (VEGFA), Fibroblast Growth Factor 1 (FGF1), and Ephrin B2 (EFNB2) for podocytes, which is consistent with their well-known roles in this compartment (**Supplementary Fig. 20; Supplementary Table 15**). We also investigated whether ligands appeared to be shared between the UB and NPC distal populations, as AVP seemed to be, finding that Neuroblastoma overexpressed (NOV) and midkine (MDK) signals promote distal tubule/connecting tubule fates across both lineages (**Supplementary Fig. 21; Supplementary Table 18**).

It is important to note that while CellRank identified some of the cell differentiation-driving interactions, many interactions with a high correlation to differentiation processes—such as IGF2, AVP, and AGT—were not identified by CellRank (**Supplementary Tables 15–17**). This may be explained by the low expression of ligands in scRNA-seq data.

In summary, we generated new, unbiased ligand-receptor interaction maps for the developing kidney. We show that these interactions correlate tightly with cellular differentiation trajectories, enabling us to identify candidate cell-cell interactions that mediate specific cell fate choices—including our newly defined cell fate convergence of NPC cells (PT transitioning to PEC and podocytes) and the convergence of NPC and UB cells (connecting tubule segment). Our data indicate the key role of ligand-receptor interactions in the spatial awareness of cells.

### Novel unbiased spatial neighborhoods-based microenvironments in developing human kidney

While we were able to show that discrete morphologic structures (blastema, PES, etc.) demonstrated a clear progression of expression and differentiation status toward mature cell types, we recognize that these structures are heterogenous and exist along a continuum. Therefore, we sought to develop an unbiased approach to characterize development based on aggregate microenvironmental effects, which we approximate using all of the calculated ligand scores. This neighborhood identification differs from our previous niche calling (and all current approaches) that focused on cell type annotations within a fixed distance.^54^

We used the ligand environment matrix, instead of the conventionally used gene expression, to generate a UMAP. This UMAP showed that specific marker signals and cell fate choices localized to specific regions within the UMAP, allowing us to cluster these neighborhoods (**Fig. 6a**). We then used this UMAP to identify neighborhoods (**Fig. 6a; Supplementary Fig. 22b, c**). In this analysis each cluster was defined by ligand abundance (**Supplementary Table 16**). For example, we identified GDNF neighborhoods (likely blastema, NPC), VEGFA neighborhood (likely glomeruli) and AVP ligand/receptor neighborhoods (likely collecting duct). These ligand/receptor neighborhoods also correlated with cell renewal of differentiation probabilities.

**Figure 6.**
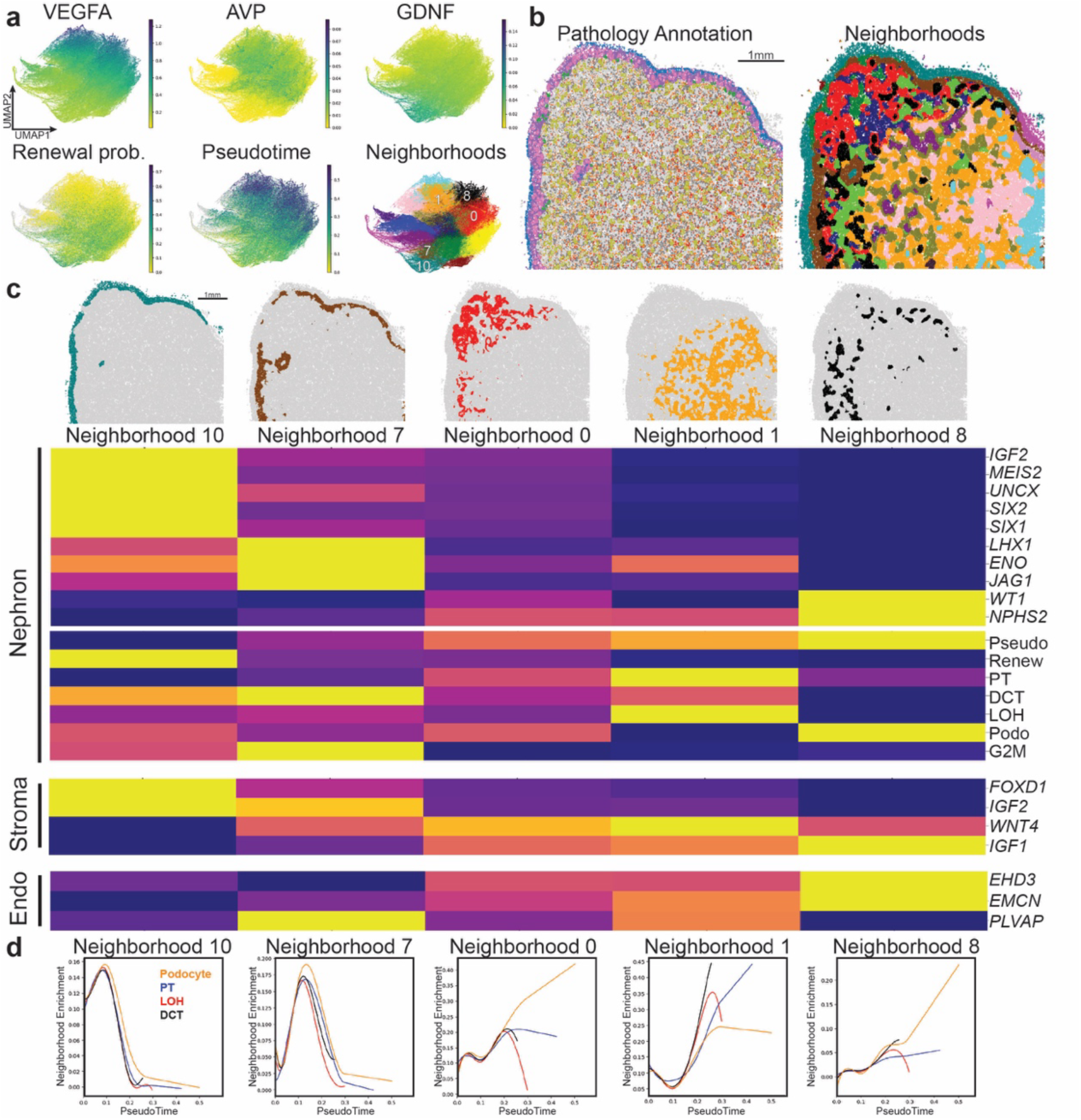
Novel unbiased spatial neighborhoods-based microenvironments in developing human kidney. (**a**) UMAPs of neighborhood signals. Color of each cell indicates signal presence (top), or absorption probability (lower left) or neighborhood (lower right). For signal presence/absorption probability low values are indicated as yellow and high being green. Neighborhoods are colored as follows: Neighborhoods: 0: red, 1: orange, 2: yellow, 3: green, 4: blue, 5: purple, 6: pink, 7: brown, 8:black, 9:dark blue, 10: turquoise, 11: gold, 12: light green, 13:cyan, 14: light purple, 15: maroon, 16: dark purple, 17: dark pink, 18: dark yellow, 19: silver, 20: dark brown, 21: reddish orange, 22: dark orange (also shown in Supplementary Fig. 20c). (**b**) Histologically defined unbiased neighborhoods in space (left) and microenvironment defined neighborhoods in space (right). Neighborhoods are indicated by color. (**c**) The heatmap of gene expression and trajectory characteristics of microenvironment derived neighborhoods. Unbiased neighborhood spatial location are shown above and the columns under show representative gene expression of heatmap. Gene expression and trajectory characteristics are rows. Low values are indicated in purple and high values by yellow. (**d**) Pseudotime trajectory characteristics of each microenviroment derived neighborhood. For each microenvironemnt neighborhood, we graph aggregate trajectory enrichment (y-axis) against pseudotime (x-axis) for each linage, as indicated by color. PT: proximal tubule, LOH: loop of Henle, DCT: distal convoluted tubule.

To verify that these ligand-based neighborhoods captured relevant biology, we compared them with our histologic neighborhoods (**Fig. 6b**). We found that microenvironment signal-based neighborhoods shared similarities with the histologically defined neighborhoods, differentiating the blastema from other portions of the nephrogenic zone, the nephrogenic zone from the cortex, and identifying medullary regions (**Fig. 6b; Supplementary Fig. 22d**). Most importantly, this approach identified additional spatially defined developmental environments, indicating the existence of further additional developmental stages not fully visible for the pathologist. We found that these neighborhoods predicted gene expression for nephron, endothelial, and stromal cells, as well as nephron progenitor cell (NPC) lineage trajectories (**Fig. 6c**). In examining the absorption probability per neighborhood, we found that neighborhood 6, which spatially resembles the blastema, corresponded with NPC renewal; neighborhood 5 with early NPC differentiation; neighborhood 1 with early tubular differentiation; neighborhood 0 with late tubular differentiation; and neighborhood 7 with renal corpuscle maturation (**Fig. 6c, d**). We noted that these neighborhoods also correlated with the proximal tubule (PT) fate (**Supplementary Fig. 22e**). Specifically, we found that PT cells with a high renal corpuscle absorption probability show markers of normal PT cells based on both raw CosMx counts and imputed gene expression, demonstrating that these cells are indeed PTs and not intermediate cell fates (**Supplementary Fig. 22f**). These results further confirmed our new NPC differentiation model and the observed plasticity.

Overall, we show that the cell-cell sender signal matrix identified new biologically meaningful cellular neighborhoods definition, which redefines human kidney development. We also validate IGF2 as a potential renewal factor.

## Discussion

We present a comprehensive analysis of human fetal kidney development, driven by a pioneering combination of the first spatially resolved single-cell gene expression atlas, high-quality scRNAseq, and advanced computational analyses. Using a probabilistic cell trajectory analysis we improve the understanding of cell differentiation trajectories over kidney development, including an early bifurcation between renal corpuscle and tubular lineages. Notably, we identify cellular plasticity, showing that already differentiated proximal tubule cells transition into parietal epithelial cells (PECs) and podocytes. Crucially, our spatial transcriptomic atlas not only captures the spatial dimension of cellular differentiation but also enables us to map the specific anatomical regions where differentiation trajectories diverge. This offers a novel perspective on the physical context of key developmental decisions. Additionally, through ligand-receptor interaction analysis, we provide a refined understanding of developmental structures and offer the first insights into how local signaling environments influence early cell fate decisions and later cell fate convergence. We identify critical ligands that govern these processes, offering new avenues for therapeutic exploration in kidney development and disease.

In this study, we improve the accuracy of cellular differentiation trajectories in the human kidney. Previous research in mouse kidney development indicated a sequence of binary decisions: First a choice between nephron progenitor cell (NPC) renewal and differentiation. Differentiating intermediates are then faced with a fate decision between pre-renal corpuscle and pretubular cell fates. Tubular cell fates are then specified by proximal or distal tubule specification, and finally to the differentiation of loop of Henle (LOH) or the distal convoluted tubule (DCT) cells. In contrast, early single-cell trajectory studies in the human kidney suggested a critical proximal-distal decision in NPC-derived intermediates, where proximal cells gave rise to both podocytes and proximal tubule (PT) cells. By employing CellRank to generate a probabilistic model of cell differentiation, our data reveal an essential renal corpuscle versus tubular fate decision at the intermediate stage, aligning more closely with the mouse nephrogenesis model. Remarkably, this first fate decision appears to be plastic, finding that PT cells appear to transition into a renal corpuscle fate becoming PECs, which can continue to transition into podocytes.^11^ This PT-to-PEC transition has been a subject of considerable debate in studies of the adult mouse kidney and has not been demonstrated previously in mice or humans, underscoring the significance of PECs as an important transitional state. Overall, our revised model highlights both an early cell fate decision between renal corpuscle and tubular fates and a late-stage cell fate convergence, offering new insights into kidney development.

In this study, we provide a comprehensive, unbiased spatial characterization of the developing human kidney gene expression at single-cell resolution. By integrating whole-transcriptome scRNA-seq and single-cell spatial transcriptomic data, we generated a single-cell resolution map of the developing human kidney. This map enabled us to align developmental stages and trajectories onto histological slides. Our integrated single-cell spatial gene expression and cell trajectory analysis strongly correlated with classic developmental structures observed on histological slides. We offer a detailed gene expression, cell interaction, and developmental trajectory map for these classic developmental programs in human kidneys. Notably, we observed that traditional histological structures do not correspond to discrete transcriptional states, as evidenced by continuous intermediate cells on the UMAP. Therefore, we developed an unbiased approach to characterize development based on aggregate microenvironmental effects, using all identified microenvironmental signals to define spatial neighborhoods. This neighborhood identification differs from previous niche calling methods that focus on cell type annotations within a fixed distance.

Ligand-receptor interactions play a key role in enabling cells to interpret spatial cues, which is crucial for spatially and temporally coordinated differentiation to form complex organs. Using our ability to define cell differentiation trajectories in space, we aimed to identify specific cell-cell interactions that regulate these trajectories in an unbiased manner. Traditional ligand-receptor interaction analyses are limited in scRNA-seq by the lack of spatial proximity information. Our new spatial gene expression data enabled a spatially aware ligand-receptor analysis, with full-transcriptome estimated expression permitting widespread discovery. We generated new, unbiased ligand-receptor interaction maps for the developing kidney and found that these interactions correlate tightly with cellular differentiation trajectories. This enabled us to identify candidate cell-cell interactions that mediate specific cell fate choices—including our newly defined cell fate convergence of nephron progenitor cells (NPCs) transitioning from proximal tubule (PT) cells to parietal epithelial cells (PECs) and podocytes, as well as the convergence of NPC and ureteric bud (UB) cells in the connecting tubule segment. Overall, we show that the ligand-receptor interaction matrix can identify new biologically meaningful cellular neighborhood definitions for human kidney development. We also identified Insulin-Like Growth Factor 2 (IGF2) as a potential renewal factor, highlighting the key role of ligand-receptor interactions in the spatial awareness of cells. Interestingly, IGF signaling is known to be critical to the development of the pancreas, though this is thought to be primarily driven by IGF1, not IGF2.^55^ The extent to which this biology is conserved between organs may be intriguing for future study.

We believe that our approach to dissecting spatially driven signals in human development has broad significance for understanding how the microenvironment affects cellular differentiation across developmental systems. While the role of cellular neighbors and ligands has long been acknowledged, our method permits broad *a priori* discovery by examining the effects of neighboring cells in greater detail. Identifying candidate ligands that drive cellular differentiation may improve systems for differentiating stem cells *in vitro* into organoids or even whole organs. Furthermore, we suspect that interpreting candidate signals is less likely to be complicated by complex feedback loops than analyzing gene expression alone. In addition, we developed a microenvironment-based neighborhood definition that differs from previous methods, which have largely focused on the presence of neighboring cell types within a certain distance to perform clustering. Currently, our approach is based on ligand-receptor interactions, but it could be further enhanced by including factors such as the extracellular matrix components, physical forces (mechanical stress), and chemical factors (e.g. pH, metabolites). Overall, our data underscore the critical role of microenvironments in cell fate determination and plasticity. We identify key ligands that likely mediate the convergence of cell types during human kidney development, such as the transitions between proximal tubule (PT) cells, parietal epithelial cells (PECs), podocytes, ureteric bud (UB) cells, and nephron progenitor cells (NPCs).

In summary, we not only provide a revised understanding of human kidney differentiation but also present the first comprehensive analysis of how microenvironmental signals affect organogenesis. We achieved this by integrating single-cell RNA sequencing (scRNAseq) with a single-cell spatial transcriptomics atlas of the human fetal kidney. Our findings reveal that the diverse microenvironments of the kidney are highly spatially organized and closely associated with diverging and converging developmental trajectories. We identified individual ligand/neighboring cell signals that predict cell fate decisions, allowing us to demonstrate the human relevance of mechanisms previously examined in mice using genetic tools, as well as identifying potential novel key mediators of cell fate across lineages.

### Limitation of the Study

Due to technical limitations in spatial gene expression data, we mapped scRNAseq counts onto our CosMx dataset to enable full transcriptome mapping in space, rather than relying solely on measured counts. This approach helps us avoid issues such as transcript dropout, a limited spatial probe library, and segmentation errors (e.g., RNA counts being misattributed to neighboring cells), but it relies heavily on the quality of the integration. To minimize integration errors, we removed CosMx cells that lacked high similarity to scRNAseq cells, which may slightly limit the scope of our atlas.

Furthermore, our estimation of the microenvironment using CytoSignal signal scores relies on pre-reported interactions (878 in total). While this is a substantial dataset, it does not capture all relevant interactions, particularly those specific to mechano-transduction. Most importantly, individual candidate ligands and transitions will require biological validation, but the number of newly identified driver interactions is beyond the scope of the current manuscript.

## Acknowledgements

Work was supported by 2021164 Bilateral US Israel grant to Drs. Susztak and Pleniceanu. JL is supported by an ASN Ben Lipps Fellowship. BD is supported by the Deutsche Forschungsgemeinschaft (DFG, German Research Foundation), Grant ID: DU 2449/1-1. This work was supported partly by Open Philanthropy projects (10080664 and 10095457) to K. Sasaki. We would also like to thank Mark Knepper, Louis Prahl for their advice.

## Author Contributions

JL, SG, AA, BD, AH, JF, ZM, RLK, BW, GR and AMB performed experiments. JL and BD performed computational analysis. BD, RLK, EH, KAK, OP, ML, PT, AJH, NZ, JP, PB and K. Sasaki offered experimental and analytical suggestions. K. Susztak was responsible for overall design and oversight of the experiments. LF performed pathological scorings. K. Susztak supervised the experiment. JL and K. Susztak wrote the original draft. All authors contributed to and approved the final version of the manuscript.

## Supplementary Materials

**Supplementary Materials 1: 3D-UMAP of nephron cell states.** 3D dimensional UMAP from scRNAseq data of nephron cell states demonstrating PECs being a potential intermediate population between PT cells and podocyte populations.

## Supplementary Tables

**Supplementary Table 1: Sample Characteristics for scRNAseq and CosMx samples.** Gestational age, additional clinical history and total number of cells assayed for 5 scRNAseq and 3 CosMx samples.

**Supplementary Table 2: scRNAseq markers for annotated cell types.** Wilcoxon test was performed, and the top 100 most statistically significant genes are show for each annotated cell type population in our scRNAseq dataset.

**Supplementary Table 3: Age Dependent Gene comparison for NPCs.** Genes associated with gestational age were calculated across the scRNAseq data for pseudobulked NPC cell type annotations. Higher logFC indicates increasing expression with gestational age.

**Supplementary Table 4: Age Dependent Gene comparison for PT cells.** Genes associated with gestational age were calculated across the scRNAseq data for pseudobulked PT cell annotations. Higher logFC indicates increasing expression with gestational age.

**Supplementary Table 5: Age Dependent Gene comparison for podocyte cells.** Genes associated with gestational age were calculated across the scRNAseq data for pseudobulked podocyte cell annotations. Higher logFC indicates increasing expression with gestational age.

**Supplementary Table 6: Age Dependent Gene comparison for LOH cells.** Genes associated with Agestational age were calculated across the scRNAseq data for pseudobulked LOH cell annotations. Higher logFC indicates increasing expression with gestational age.

**Supplementary Table 7: Age Dependent Gene comparison for DCT cells.** Genes associated with gestational age were calculated across the scRNAseq data for pseudobulked DCT cell annotations. Higher logFC indicates increasing expression with gestational age.

**SupplementaryTable 8: Age Dependent Gene comparison for stromal cells.** Genes associated with gestational age were calculated across the scRNAseq data for pseudobulked stromal cell annotations. Higher logFC indicates increasing expression with gestational age.

**Supplementary Table 9: Age Dependent Gene comparison for endothelial cells.** Genes associated with gestational age were calculated across the scRNAseq data for pseudobulked endothelial cell annotations. Higher logFC indicates increasing expression with gestational age.

**Supplementary Table 10: Gene drivers for NPC renewal.** Correlations between gene expression and the CellRank absorption probabilities for NPC and an aggregate endpoint of any terminal cell type (PT, LOH, DCT or podocyte).

**Supplementary Table 11: Gene drivers for tubular and renal corpuscle specification.** Correlations between gene expression and the CellRank absorption probabilities for tubular fates (aggregate of LOH, PT and DCT cell states) and renal corpuscle cell fates (podocytes).

**Supplementary Table 12: Gene drivers for terminal cell.** Correlations between gene expression and the CellRank absorption probabilities for each terminal cell fate (PT, LOH, DCT, podocyte).

**Supplementary Table 13: Cell-cell interactions as estimated by CellPhonedb using histologic neighborhood and mature cell type.** Using the histologic classifications for nephrogenic zone cells and cell type annotation for cells outside of the nephrogenic zone, CellPhonedb scores are shown.

**Supplementary Table 14: Cell-cell interactions as CytoSignal.** Ligand-receptor interactions were called for each sample. Interactions conserved across all 3 samples are also displayed.

**Supplementary Table 15: Microenvironment signal correlation with absorption probabilities.** Environmental cue scores from CytoSignal and their correlation with absorption probabilities for renewal, differentiation, PT, podocyte, LOH, DCT, tubule, renal corpuscle cell fates. Pseudotime correlations with each ligand is also shown.

**Supplementary Table 16: Differentially abundant ligands within each neighborhood.** Differentially present ligands are shown as markers for each of the environmentally determined neighborhoods.

**Supplementary Table 17: Microenvironment signal correlation with absorption probabilities of tubular and renal corpuscle fates for PT cells.** Environmental cue scores from CytoSignal and their correlation with absorption probabilities for tubule, renal corpuscle cell fates within the PT population.

**Supplementary Table 18: Microenvironment signal correlation with absorption probabilities of UB lineages.** Environmental cue scores from CytoSignal and their correlation with absorption probabilities for connecting tubule (CNT), urothelium and intercalated cells (IC).

## Materials and Methods

### Fetal Sample Acquisition

Human fetal kidneys were obtained from donors undergoing elective abortion at the Gynecology Clinic, University of Pennsylvania, Philadelphia and from the Sheba Medical Centre, Tel Aviv, Israel. Donors provided informed consent. All experimental procedures were approved by the Institutional Review Boards at the University of Pennsylvania (#832470) and at Sheba (approval number 8993-21S-MC). Embryo ages were determined through ultrasonographic measurement of crown rump length. Kidney tissue was dissected in RPMI-1640 medium (Gibco) using a dissection microscope.

### Mouse explants

Embryos dissected from euthanized female CD1 mice (Charles River Production) were used at e13 developmental stages for explant experiments involving IGF2. For linsitinib experiments, C57BL/6 mice were used for availability purposes. Animal studies were approved by the University of Pennsylvania IACUC.

### scRNAseq

Fresh human kidneys were collected in RPMI media and using a razor blade were cut into small pieces 2-4 mm in diameter. Tissue was furthered dissociated with gentleMACS C tube (Miltenyi, #130-110-201) using program Multi_B. Tissue was digested with Enzyme D, Enzyme R and Enzyme A in RPMI at 37C for 30 minutes. Tissue was homogenized again with gentleMacs using the Multi_B program was then performed. The resultant solution was strained through a 70 um filter, and then centrifuged at 600g for 7 minutes. The cell pellet was incubated in RBC lysis buffer for 3 minutes on ice, followed by addition of 10 mL of PBS and centrifuged again for 5 minutes at 500g. Supernatant was removed, and the pellet was resuspended in PBS. Cell number and cell viability was examined using the Countess AutoCounter (Invitrogen, C10227), and manual examination. An estimated 20,000 cells, based on AutoCounter results, were loaded into the Chromium Controller (10X Genomics, PN-120223) in a Chromium Next GEM chip G (10X Genomics, Single Cell Kit PN-1000120). The cDNA and gene expression library were made using the Chromium Next GEM Single Cell 3′ GEM Kit v3.1 (10X Genomics, PN-1000121) and Single Index Kit T Set A (10X Genomics, PN-120262) according to the manufacturer’s protocol. Quality control for the libraries was performed using Agilent Bioanalyzer High Sensitivity DNA kit (Agilent Technologies, 5067-4626). Libraries were sequenced on Illumina Novaseq 6000 system with 2 × 150 paired-end kits. Reads were then demultiplexed by sample.

### CosMx Sample preparation

Tissue sections were cut at 5 um thickness and prepared according to the manufacturer specifications (NanoString Technologies). The human universal cell characterization RNA probes were used with an 50 additional custom probes set for the following genes, which were manually selected: *ESRRB, SLC12A1, UMOD, CD247, SLC8A1, SNTG1, SLC12A3, TRPM6, ACSL4, SCN2A, SATB2, STOX2, EMCN, MEIS2, SEMA3A, PLVAP, NEGR1, SERPINE1, CSMD1, SLC26A7,SLC22A7, SLC4A9, SLC26A4, CREB5, HAVCR1, REN, AP1S3, LAMA3, NOS1, PAPPA2, SYNPO2, RET, LHX1, SIX2, CITED1, WNT9B, AQP2, SCNN1G, ALDH1A2, CFH, NTRK3, WT1, NPHS2, PTPRQ, CUBN, LRP2, SLC13A3, ACSM2B, SLC4A4, PARD3, XIST,UTY*. We used the DAPI, CD298/B2M, CK8/18, and PanCK/CD45 for additional staining per NanoString protocol. Imaging was performed using configuration A. After imaging was completed, the flowcell was incubated in 100% xylene overnight and the coverslip was removed from the slide with a razor blade. The slide was then stained with hematoxylin and eosin and imaged. The adjacent section of each tissue piece was also stained with hematoxylin and eosin and imaged as well.

### Fetal Explant culture

Fetal kidneys from e13 mice were dissected and placed on a transwell plate (#230615 Celltreat) and cultured in DMEM with penicillin/streptomycin with the addition of 500 ng/ml IGF2 (R&D, 792-MG), 70 uM Insulin (Novalog) or PBS. Media was changed daily, and after 5 days of culture, the explants were fixed in 4% PFA at room temperature for 10 minutes. Experimental comparisons were only made within the same litter of mice. Linsitinib (Selleckchem: OSI-906) experiments were performed the same way using 40, 4, 0.4 uM concentrations or DMSO being added to the media and changed daily.

### Human Nephron Organoid Generation

Nephron progenitor cell organoids were generated from the SV20 iPSC line (PENN123i-SV20, Induced Pluripotent Stem Cell Core, University of Pennsylvania, RRID:SCR_022426) according to published protocols^56–58^ iPSCs were maintained in standard tissue-culture treated 6-well plates in stem cell maintenance medium plus supplement (mTeSR+ kit, StemCell Technologies 100-0276). Cells were passaged using Accutase (StemCell Technologies, 07920) and plated at a density of 5,200 cells/cm^2^. Differentiation was induced with TeSR-E6 Medium plus supplements (TeSR-E6+) (STEMCELL Technologies 05946) and 7µM CHIR99021 (Tocris, 4423) the following day, and media was refreshed every other day for 5 days. On day 5, wells were washed with DPBS and media was changed to TeSR-E6+ with 1 µg/ml heparin (Sigma-Aldrich H4784) and 200 ng/ml FGF9 (R&D Systems 273-F9-025). Media was refreshed every day for 5 days. On day 10, cells were lifted using Accutase and excess TeSR-E6+ was used to quench digestion. Cells were pelleted by centrifuging at 300g for 3 minutes and then resuspended in TeSR-E6+ and counted. An appropriate volume of cells corresponding to 300,000 cells per organoid was then transferred to microcentrifuge tubes and centrifuged at 300g for 3 minutes. The pellet was resuspended in TeSR-E6+ with 7µM CHIR99021 to create a dense cell slurry at 3 x 10^5^ cells/µl. 1 µl aliquots of the slurry were then spotted onto 6-well 0.4 µm polyester transwell membranes (CellTreat, 230607) overlying TeSR-E6+ and 7µM CHIR 99021 ‘pulse’ media with 5-8 organoids spotted per well. Organoids were ‘pulsed’ for two hours. Following the CHIR ‘pulse’, samples were washed with DBPS before changing to chase media to TeSRE6+ with 1 µg/ml heparin, 200 ng/mL FGF9, and equivolume DPBS, 1 µM recombinant arginine vasopressin (Sigma-Aldrich, V9879), 500 ng/mL recombinant insulin growth factor 2 (R&D, 792-MG), 10 µM Tolvaptan (Sigma-Aldrich, T7455), or 10 µM recombinant angiotensin 2 (Sigma-Aldrich, A9525-1MG) for Figure (5) or 0 µM, 40 nM, 400 nM, 4 µM, or 40 µM Linsitinib (Selleckchem S1091) with equivolume DMSO for Supplementary Figure 16). Media was refreshed every day until D12 when samples were either fixed for immunofluorescence or washed with 1x DPBS before changing media to TeSRE6+ with equivolume 1x DPBS, 1 µM AVP, 500 ng/mL IGF2, 10 µM Tolvaptan, or 10 µM ANG2. Media was refreshed every other day until experimental endpoint. For “late” conditions (AVP and ANGII), organoids were cultured in PBS until day 16, at which point AGTII or AVP was added until the organoid was fixed.

### Human Nephron Organoid Immunofluorescence

At the experimental endpoint, the transwell membrane was separated from the transwell apparatus and placed in microcentrifuge tubes containing 4% paraformaldehyde for 15 minutes at room temperature, then washed twice with DPBS with 100 mM glycine, and finally exchanged with DPBS. Organoids were permeabilized with 0.5% triton x-100 in DBPS for 30 minutes at room temperature, then blocked in 1x IF wash (DPBS + 1g/l Bovine Serum Albumin + 0.2% Triton-X-100 + 0.04% Tween-20) with 5% donkey serum for 1 hour at room temperature. Primary and secondary antibody solutions were diluted in blocking solution and samples were incubated in antibody solutions overnight at 4°C. Following primary or secondary antibody incubation, samples were washed with three exchanges of 1x IF wash solution over three hours at room temperature. After the last wash, samples were briefly exchanged with DPBS before being transferred to glass slides for imaging. Once on the slides, excess DPBS was wicked away and FocusClear (Cedarlane Labs FC-101) solution was added to cover each organoid. Two 38 um spacers (Precision Brand: 44115) were added to each slide before addition of a 140 µm thick coverslip (Globe Scientific 1415-10), which was sealed to the slide with clear nail polish.

D12 organoids were stained with rabbit anti-SIX2 (1:400, 11562-1-AP, Proteintech, RRID: AB_2189084) and goat anti-JAG1 (1:150, AF599, R&D Systems, RRID: AB_2128257). D23 organoids were stained with biotinylated LTL (1:200, B-1325-2, Vector Laboratories), rat anti-ECAD (1:200, ab11512, abcam, RRID: AB_298118), sheep anti-NPHS1 (1:200, AF4269, R&D Systems, RRID: AB_2154851), rabbit anti-SIX2 (1:200), goat anti-ITGa8 (1:200,AF4076,R&D systems,RRID: AB_2296280), or mouse anti-MEIS123 (1:200, 39795, active motif, RRID: AB_2750570). Donkey secondary antibodies were used at 1:400 dilution: DyLight 405-Streptavidin (016-470-084, Jackson ImmunoResearch). Donkey anti-Mouse – 555 (Invitrogen, A31570), Donkey anti-Goat – 405 (Invitrogen, A48257), Donkey anti-Rabbit - 647 (Invitrogen, A32795), Donkey anti-Rat (A345695), Donkey anti-sheep 647 (Invitrogen, A214448), Donkey anti-goat 647 (Invitrogen: A32849).

Samples were imaged using a Nikon Ti2-E microscope equipped with a CSU-W1 spinning disk (Yokogawa), a white light LED, laser illumination (100 mW 405, 488, and 561 nm lasers and a 75 mW 640 nm laser), a Prime 95B back-illuminated sCMOS camera (Photometrics), motorized stage, 4x/0.2 NA, 10x/0.25 NA, and 20x/0.5 NA lenses (Nikon), and a stagetop environmental enclosure (OkoLabs). For 10x images of day 33 and 34 organoids, z-projections were acquired by taking 12.5 um steps spanning the entire height of each organoid with 3×3 fields of view stitched together at 1% overlap. For 10x images of day 12 organoids, z-projections were acquired by taking 5 um steps spanning the entire height of each organoid with 3×3 fields of view stitched together at 1% overlap.

### Human Nephron Organoid Imaging Analysis

Imaged organoids were analyzed in ImageJ. Jython scripts used to quantify presence of SIX2, JAG1 for day 12 organoids, as well as ECAD, NPHS1 and LTL are available on https://github.com/jlevins2010/Fetal_atlas. Briefly, area for each marker was calculated as being above a manually set intensity threshold, specific to each marker, using the “analyze particles” command for each sample to calculate area. For nephroid segments, this was calculated as total area on all images and also as a normalized area, which normalized to total nephroid area for each organoid.

### Whole Mount Immunofluorescence

After fixation, explants were blocked in 5% donkey serum and then stained with the following primary antibodies prior to imaging, JAG1 - goat (1:200), (R&D, AF599-SP), Six2 - mouse (1:100) (proteintech, 66347-1-Ig) Six2 – rabbit (1:200) (proteintech 11562-1), LTL (1:200) (Vector Laboratories B-1325), ECAD (1:200) (abcam, ab11512), NPHS1- (R&D, AF4269) sheep (1:200), ITGA8 (1:200) (R&D, AF4076). Secondary antibodies used were the following: Donkey anti-Mouse – 555 (Invitrogen, A31570), Donkey anti-Goat – 405 (Invitrogen, A48257), Donkey anti-Rabbit - 647 (Invitrogen, A32795), Conjugated EpCAM (anti-Mo), FITC (Invitrogen, 11-5791-82). Explants were washed after incubation with primary and secondary antibodies in PBS with BSA, triton and tween. Prior to imaging, explants were cleared with 1 drop of focus clear (CelExporer, FC-101). To quantify tips, Epcam positive regions of tissue were counted if at edge of tissue and bordered by Six2 positive cells. Images were analyzed using Image J, and brightness of images was adjusted to visualize the ureteric tree.

### Slide based Immunofluorescence

Sections of tissue were deparaffinized in Xylene, rehydrated in ethanol baths, and antigen retrieval was performed with incubation in Sodium Citrate (10 mM pH 6.0 at 100C for 20 minutes. Samples were blocked in 10% BSA. Primary antibodies used were 1:50 (phospho-AKT 1 (1:50, CST#9271), WT1 (1:200) (Novus NB110-60011) and (abcam CAN-R9(IHC)-56-2), ACE2 (Thermo, goat PA5-47488 and mouse MA5-32307). Slides were then washed with PBS, and then incubated with 1:400 secondary Alexaflor 555 goat-anti rabbit (Thermo #A-21428) and Alexafor 488 donkey-anti goat (Thermo: A-11055).

### Bioinformatic Analysis: scRNA-seq

Alignment of samples was performed with cellRanger^59^ (version 7.2.0) to generate the original expression matrix used for cell type annotation and connectivity and STARSOLO^60^ (version 2.7.9a) to generate spliced, un-spliced, and ambiguous gene x cell matrices. CellBender^61^ (version 0.2.0) was used to remove ambient RNA, using the following parameters, expected cells 10000, total-droplets-included 50000, False positive rate 0.01, epochs 150 and a default learning rate. Learning curves for each sample were manually inspected to ensure local training was appropriate.

Basic scRNAseq processing was performed using scanpy^62^. Doublets were removed using scrublet (version 0.2.3). HarmonyPy^63^ (version 0.09) was used for batch correction of the scRNAseq data using default parameters. Cells were filtered to remove those with greater than 25% mitochondrial counts, fewer than 500 genes. Only genes expressed in at least 3 cells in all samples were used for further analysis. Variable genes were called using scanpy (version 1.9.1) using min_mean=0.0125, max_mean=3, min_disp=0.5, batch_key = ‘sample’. Cell cycles was regressed out of each PC, and neighbors – 15 for each cell--were called using 40 PCs. Clustering was performed using the Leiden algorithm (leidenalg version 0.8.10), with resolution 1.0. Leiden clusters DEGs, called using the Wilcoxon test, were inspected to determine cell type annotation for each cluster. UMAPs were used to visualize cells in dimension reduced space (umap-learn version 0.5.3).

RNA velocity was calculated using scVelo (version 0.2.5), suing 30 neighbors and a dynamical model. CellRank^35^ (version 1.5.1) was used for calculating absorption probabilities to calculate trajectories of the single cell data, using a mixed kernel (0.5 weight connectivity and 0.5 weight velocity). 7 macrostates were calculated and 400 cells with the top score for each microstate were called. Absorption probabilities were called for each terminal state (PT, LOH, Podocyte, DCT), for renal corpuscle vs. tubular (podocyte compared to aggregate PT, LOH and DCT), and renewal/differentiation (aggregate terminal states vs. NPC).

Age dependent DEGs were calculated using pyDESEQ2^64^ (version 0.4.4) via pseudobuking each annotation by sample and plotted in matplotlib (version 3.6.3).

### Bioinformatic Analysis: Spatial single cell transcriptomics

The expression matrix and metadata from each CosMx run was exported from the AtoMx platform and was converted to a python object using squidpy^65^. All samples were merged and preprocessed and analyzed together. Cells with fewer than 30 counts were filtered out. Counts from *MALAT1* and hemoglobin genes were removed. Count matrices were normalized, and log transformed. PCA was performed on these matrices. Neighbors were calculated using 30 PCs and 15 nearest neighbors, which were used to generate an initial UMAP. Leiden clustering was performed using a resolution of 1 and initial cell type annotation was performed. Raw counts of the initial object were used for SCVI batch correction using 5 layers to generate 30 latent variables. This model was trained for a maximum of 1000 epochs using a learning rate of 0.001. These latent variables were used to recalculate neighbors. Using 15 neighbors per cell Leiden clustering was again performed for repeat annotation. Using annotations from scRNAseq cells and he CosMx cells with the 80% closest intra-technology distance, we used SCANVI to recalculate latent variables, which were used for neighbor calling (using all 30 latent variables) and Leiden clustering 15 nearest neighbors (based on expression) were annotated for the final time. Meta data and expression was imputed for each remaining CosMx cells by averaging distance squared weighted values of the most similar scRNAseq cells. Spatial neighbors were called for 25-, 50-, 75- and 100-micron distances using squidpy. Differential expression between cell types was performed using the rank genes function of scanpy using a Wilcoxon test. Generalized additive models were constructed using pyGAM (version 0.8.0) and plotted using matplotlib (version 3.6.3).

### Pathway Analysis

GSEApy^66^ (version 1.1.2) was used using the EnrichR^67^ method was used for pathway analysis. Total expressed genes measured was used for the background list and WikiPathway_2023_Human pathway list, and input genes with an adjusted p-value of < 0.05.

### Histologic neighborhood annotation

Whole slide imaging was performed on H&E-stained images of both original CosMx slides and sequential sections. These images were aligned using QuPath, and annotations were performed by a pathologist. Spatial locations from annotations were calculated using GeoPandas and merged into the scanpy object for further analysis. Differential expression between neighborhoods was performed using the rank gene function, using a Wilcoxon test. CellPhoneDb was used to identify spatially plausible interactions, using each neighborhood as a distinct microenvironment. CytoSignal was also used for spatially aware ligand-receptor analysis, with a p-value cutoff of 0.05, read threshold of 100 and a significance threshold of 100.

Spatial neighborhoods were determined using the dge.lig matrix for each sample using CytoSignal, and ligand names were named pulled from dge.lig column names and were converted from UniProt to gene symbol. These were imported into an anndata object for Leiden clustering and UMAP construction. Differential ligand expression between neighborhoods was performed using the rank genes function of scanpy using a Wilcoxon test. Generalized additive models were constructed using pyGAM (version 0.8.0) and plotted using matplotlib (version 3.6.3).

### Statistics & Reproducibility

Independent sample two-sided t-test was used to compare the likelihood that two groups were drawn from the same population using scipy stats package. A *p* < 0.05 was considered as a significant.

### Data Availability

Raw data, processed data, and metadata from the scRNAseq and CosMx spatial transcriptomics has been deposited in Gene Expression Omnibus (GEO) with the accession code of ***, reviewer token:***. Scripts have been shared at https://github.com/jlevins2010/Fetal_atlas.

