## Supplementary figures and images for "Single-Cell Spatial Mapping of Human Kidney Development Reveals the Critical Role of the Local Microenvironment in Cell Fate Decisions"

### Supplementary Fig 2

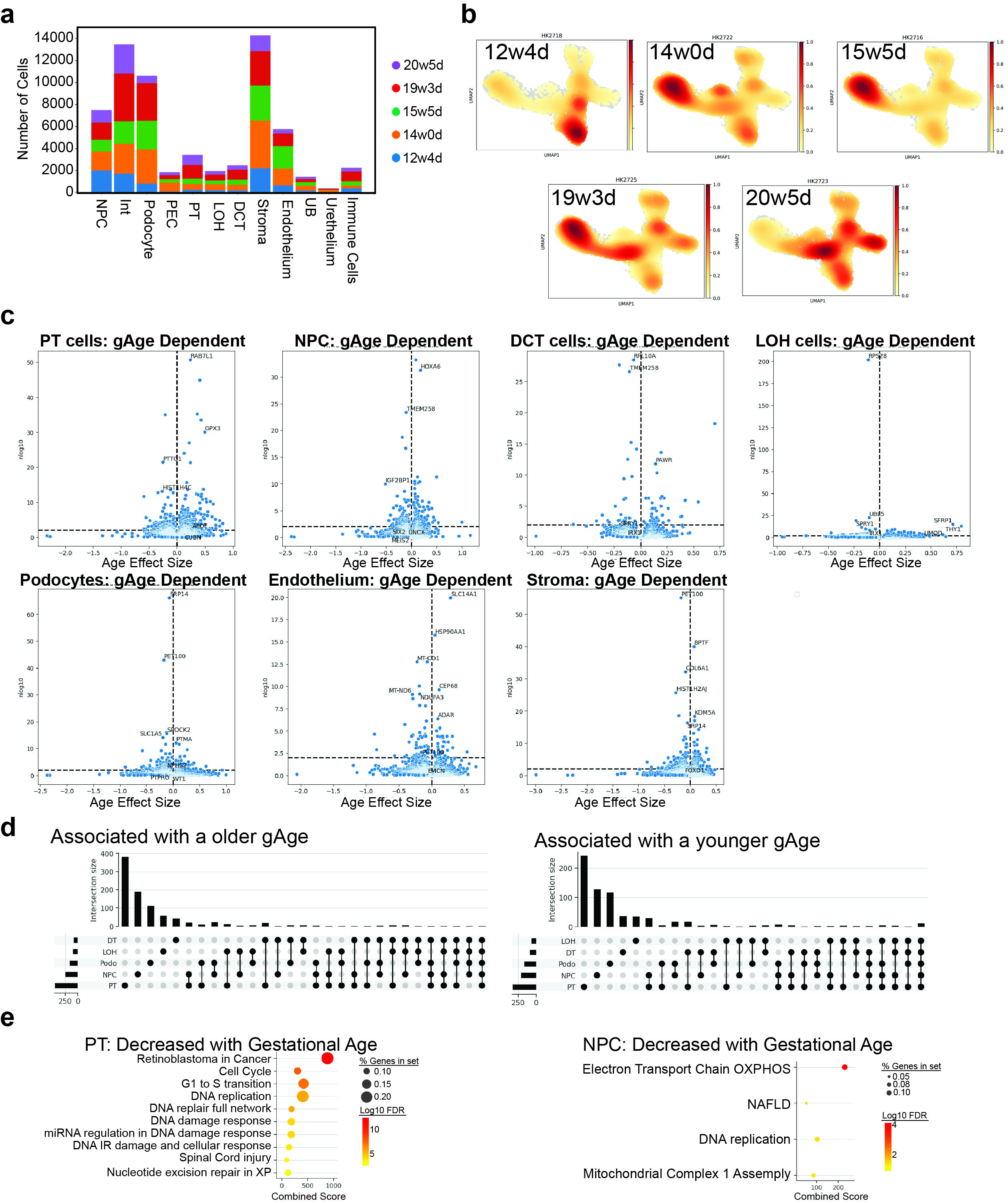

### Supplementary Fig. 1

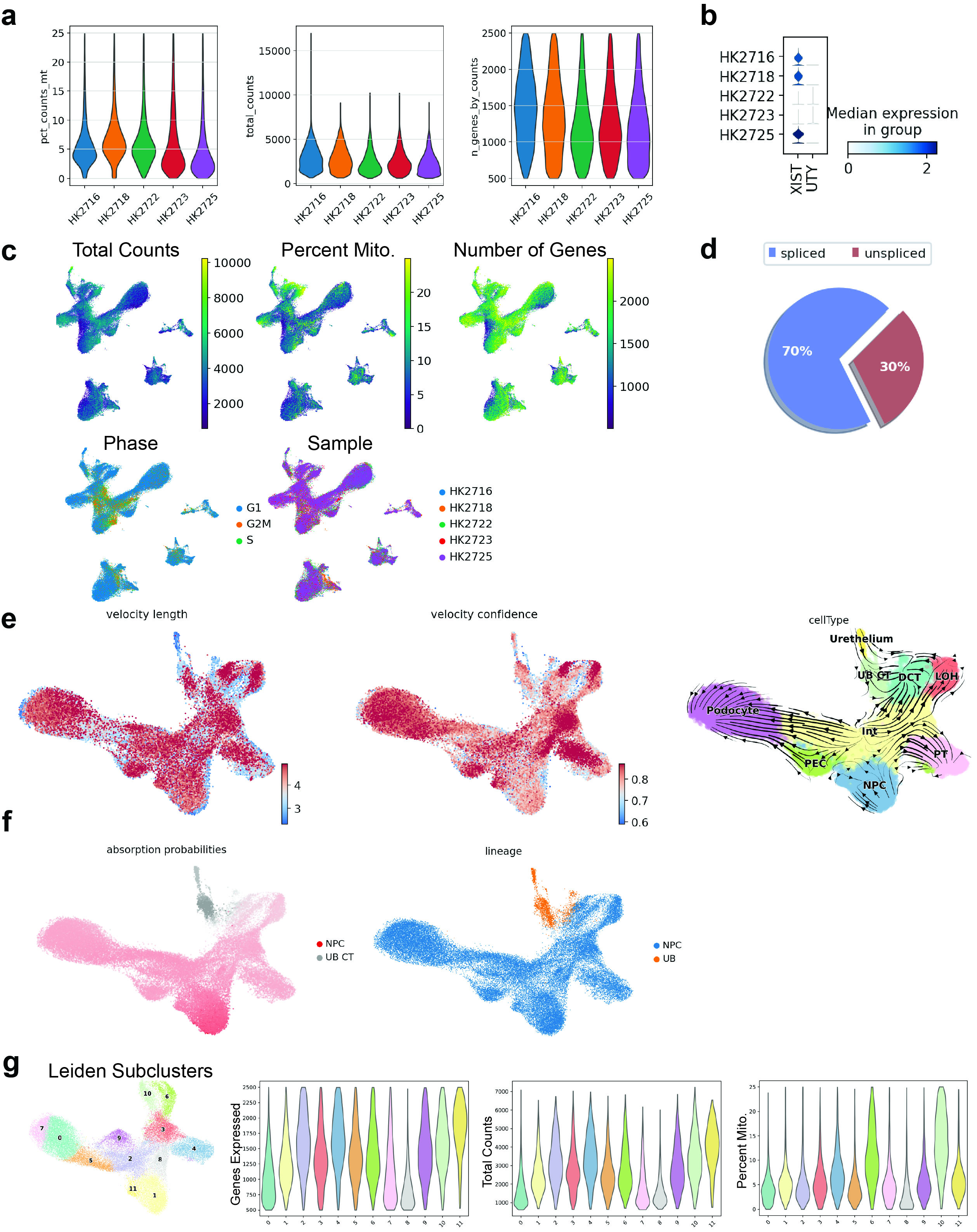

### Supplementary Fig. 3

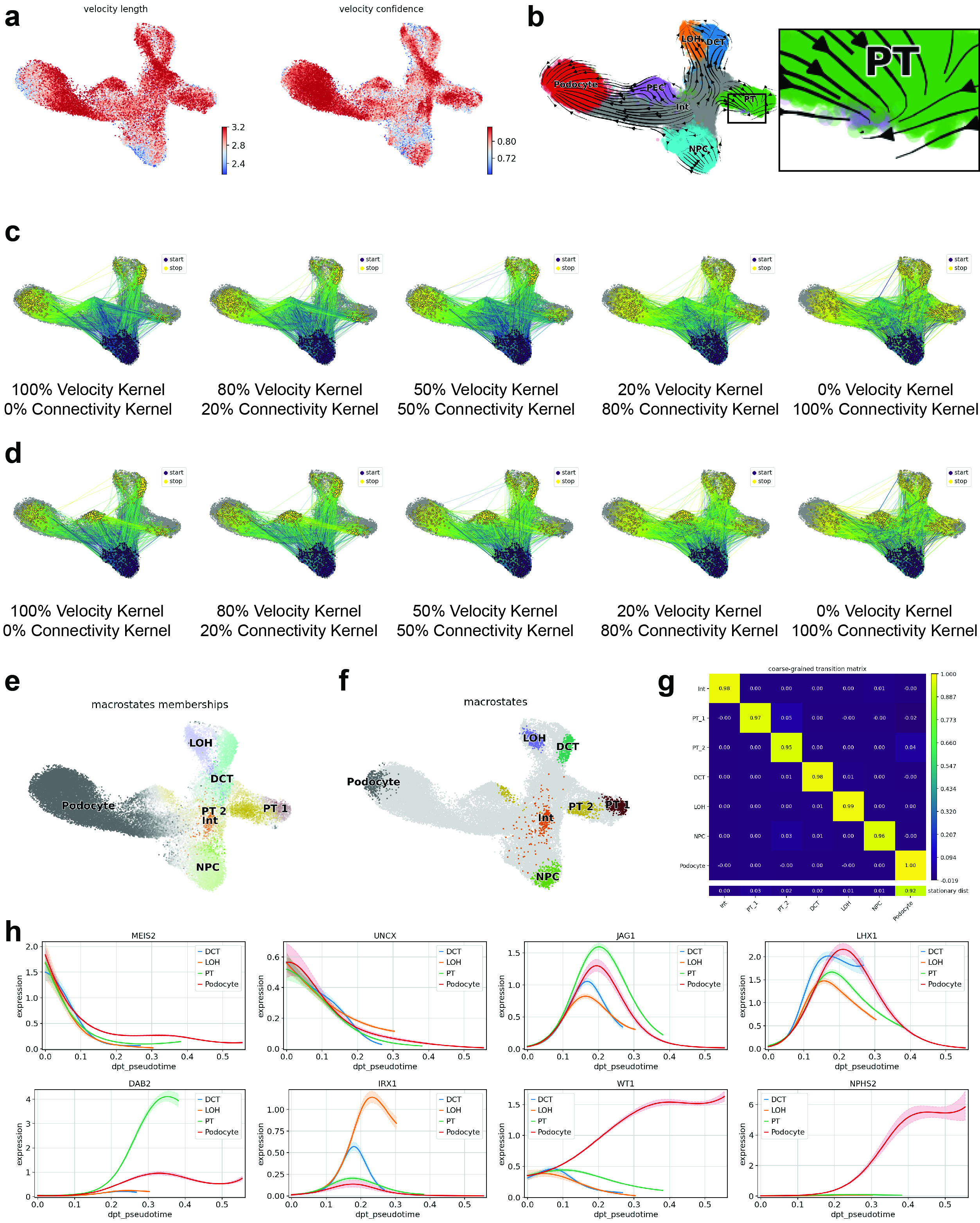

### Supplementary Fig. 4

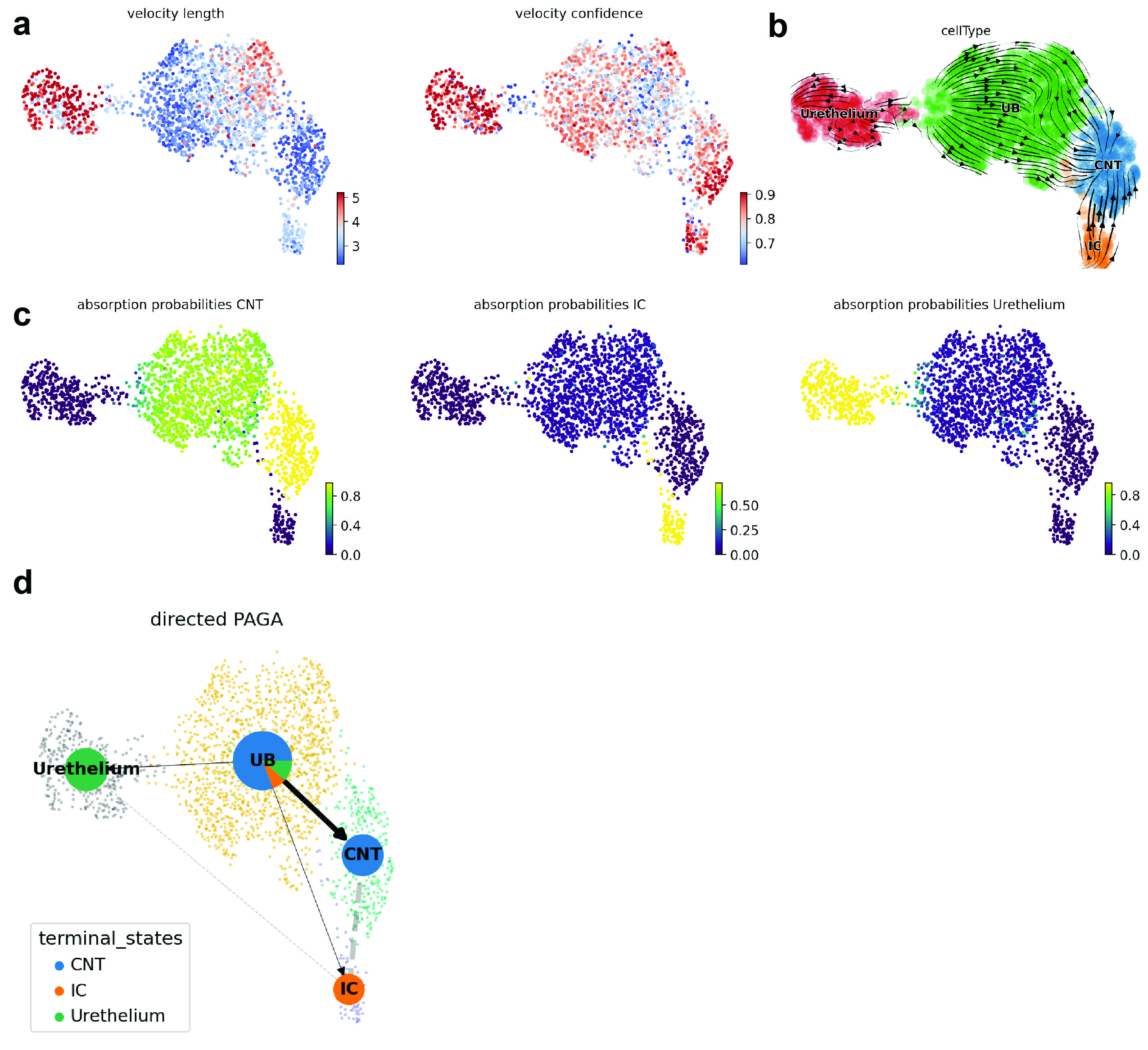

### Supplementary Fig. 5

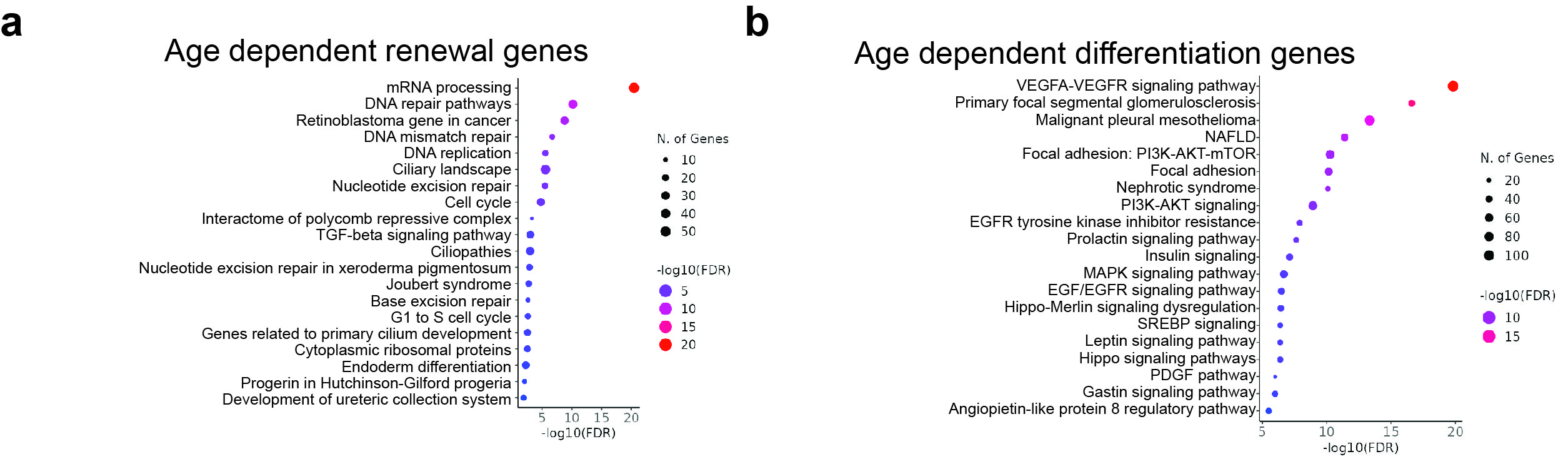

### Supplementary Fig. 6

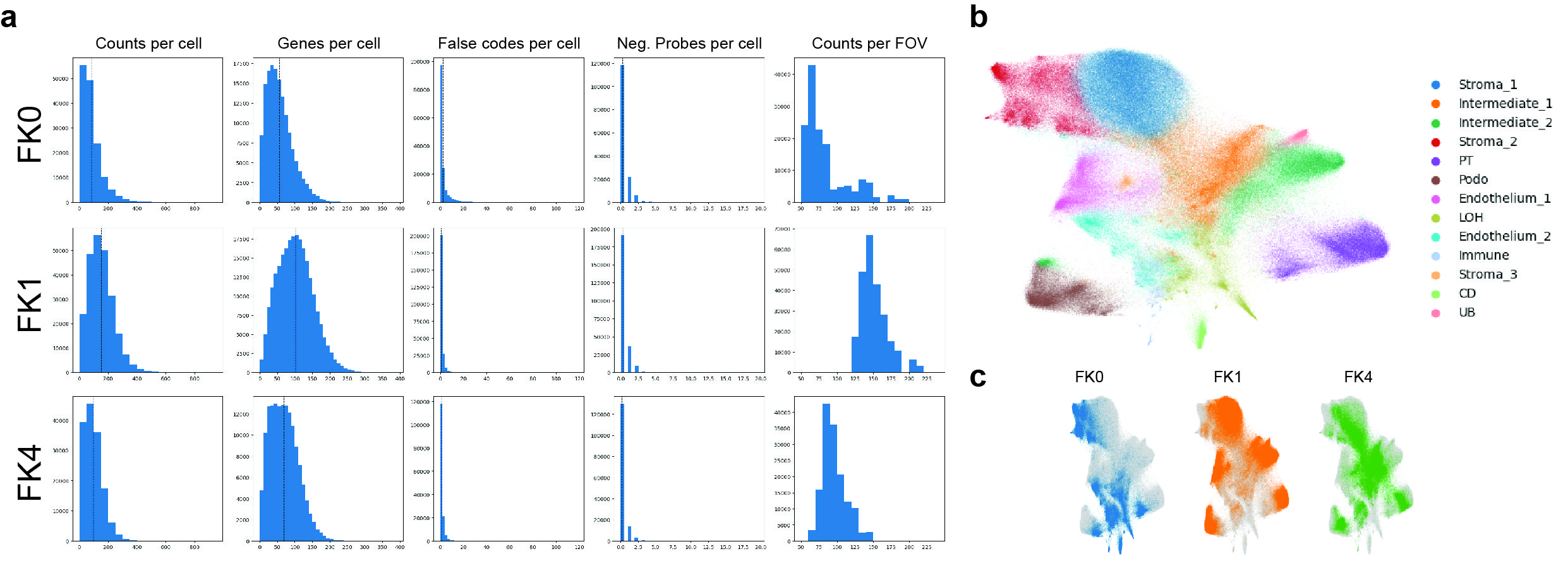

### Supplementary Fig. 7

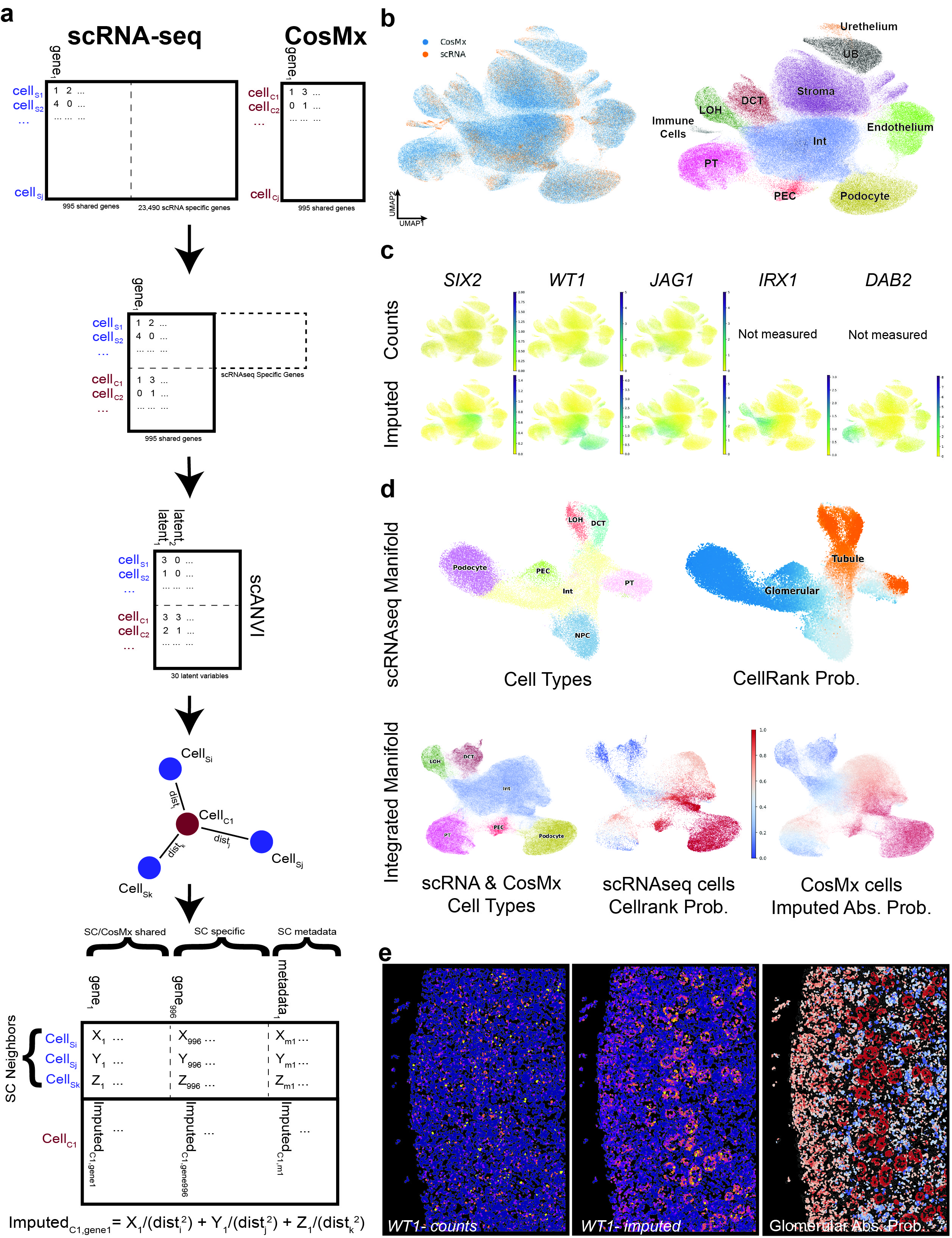

### Supplementary Fig. 8

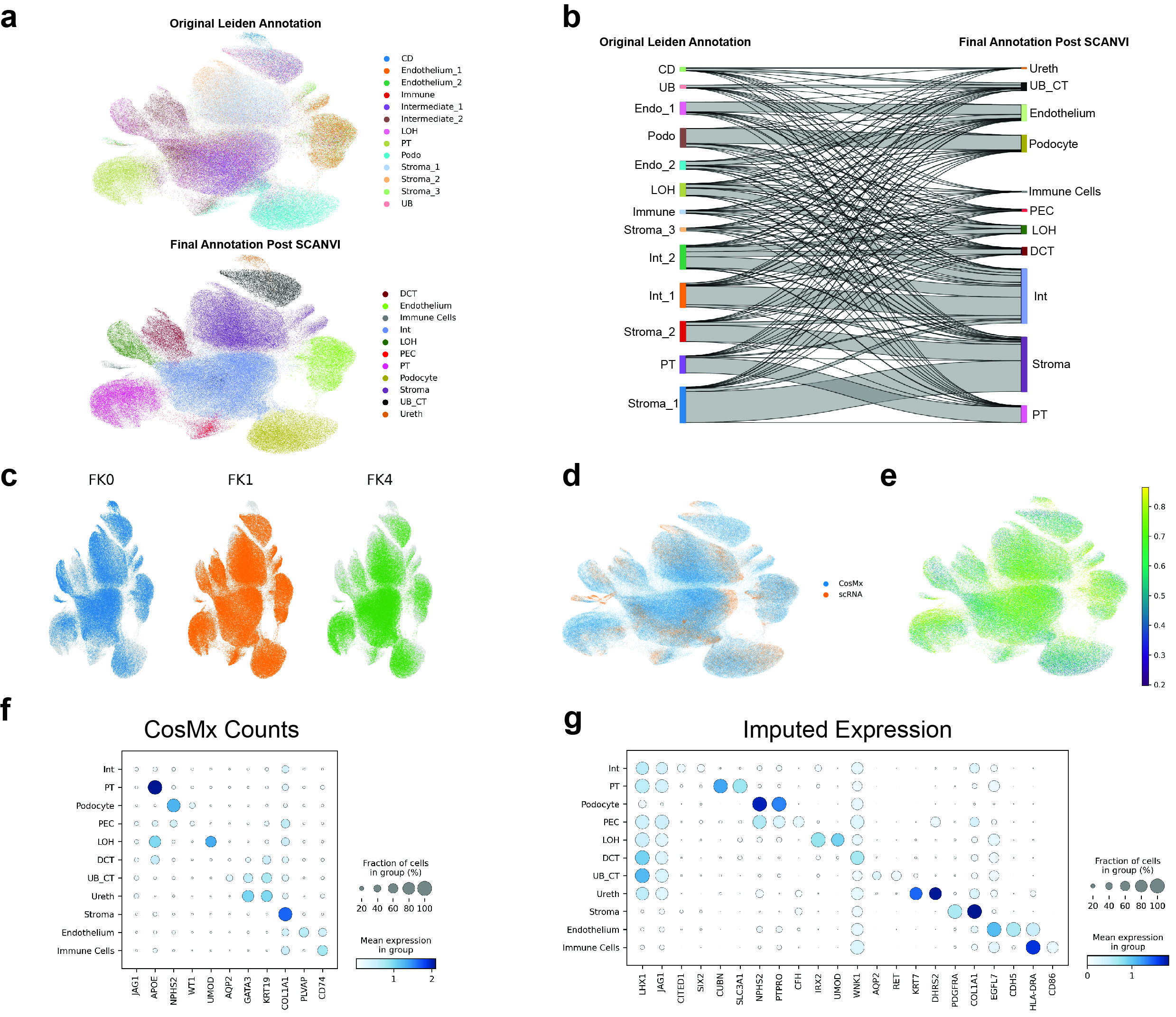

### Supplementary Fig. 9

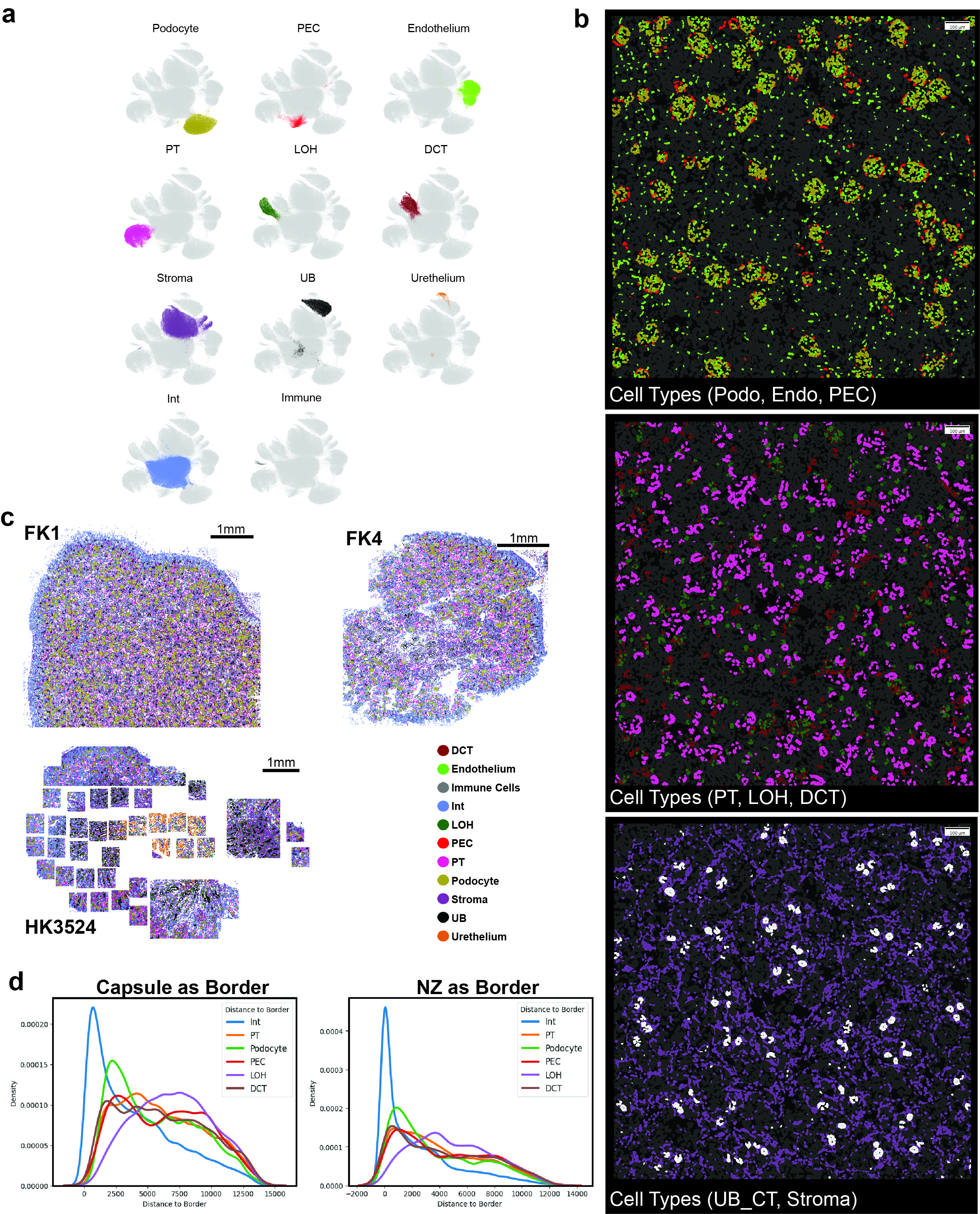

### Supplementary Fig. 10

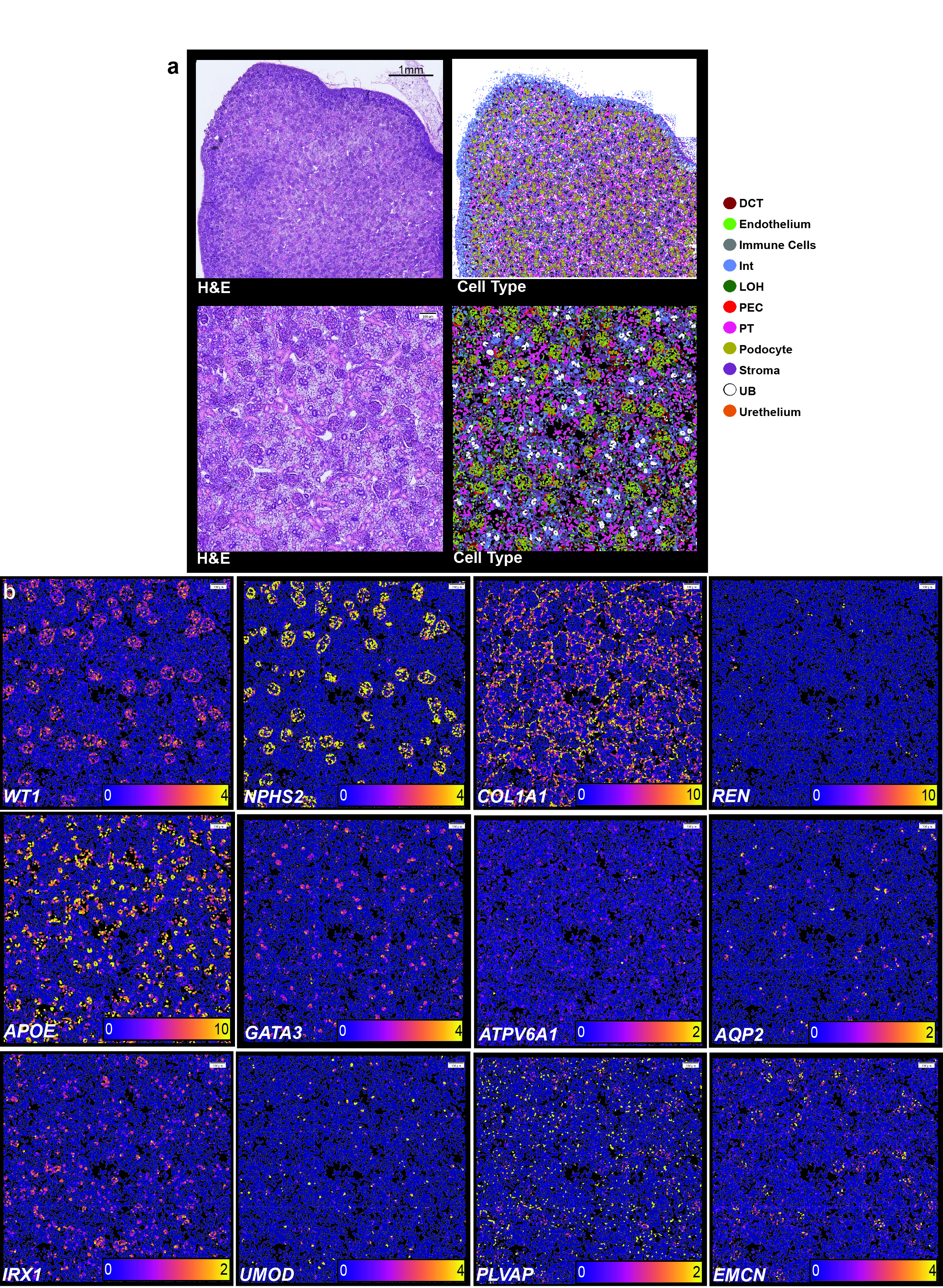

### Supplementary Fig. 11

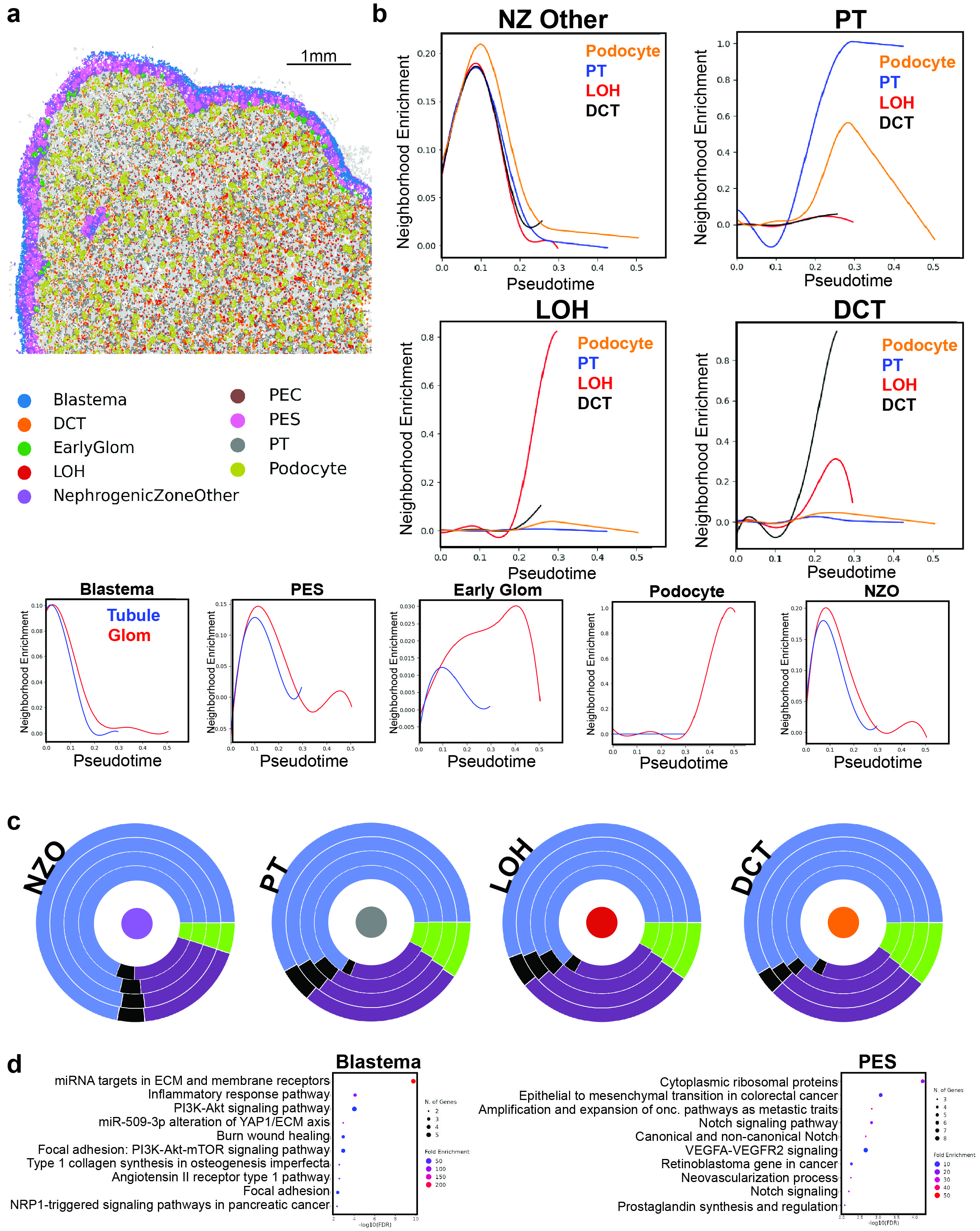

### Supplementary Fig. 12

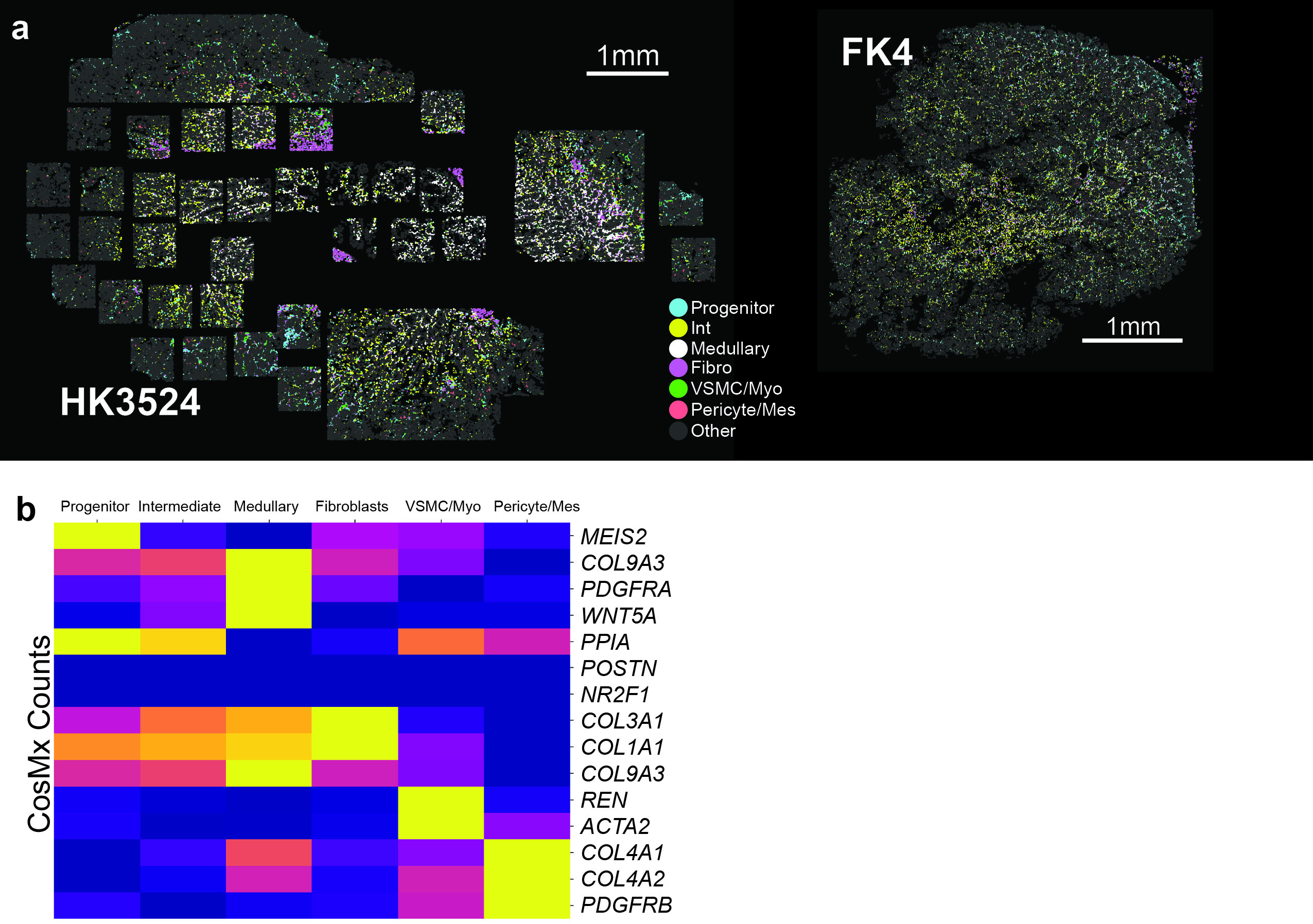

### Supplementary Fig. 13

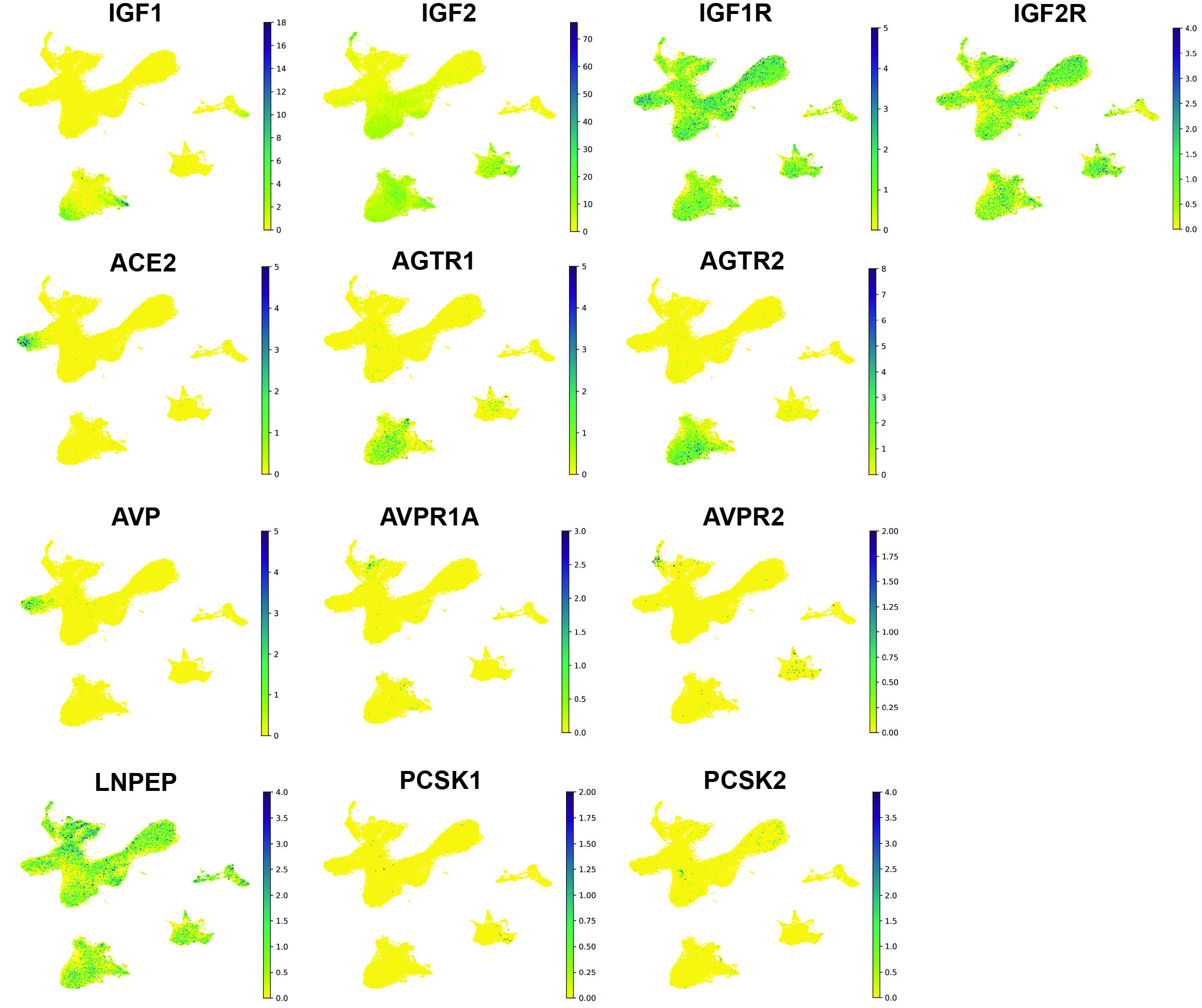

### Supplementary Fig. 13

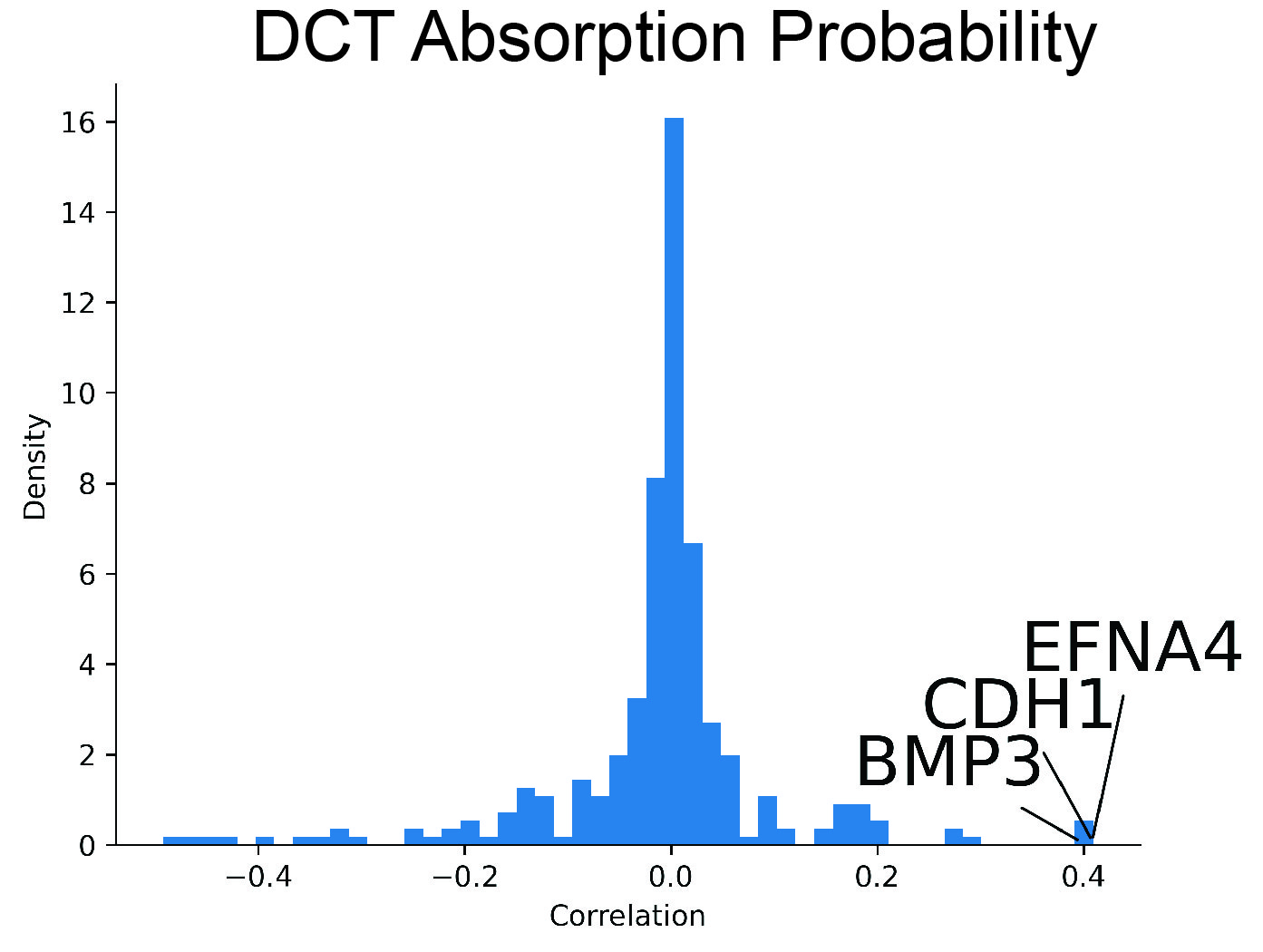

### Supplementary Fig. 15

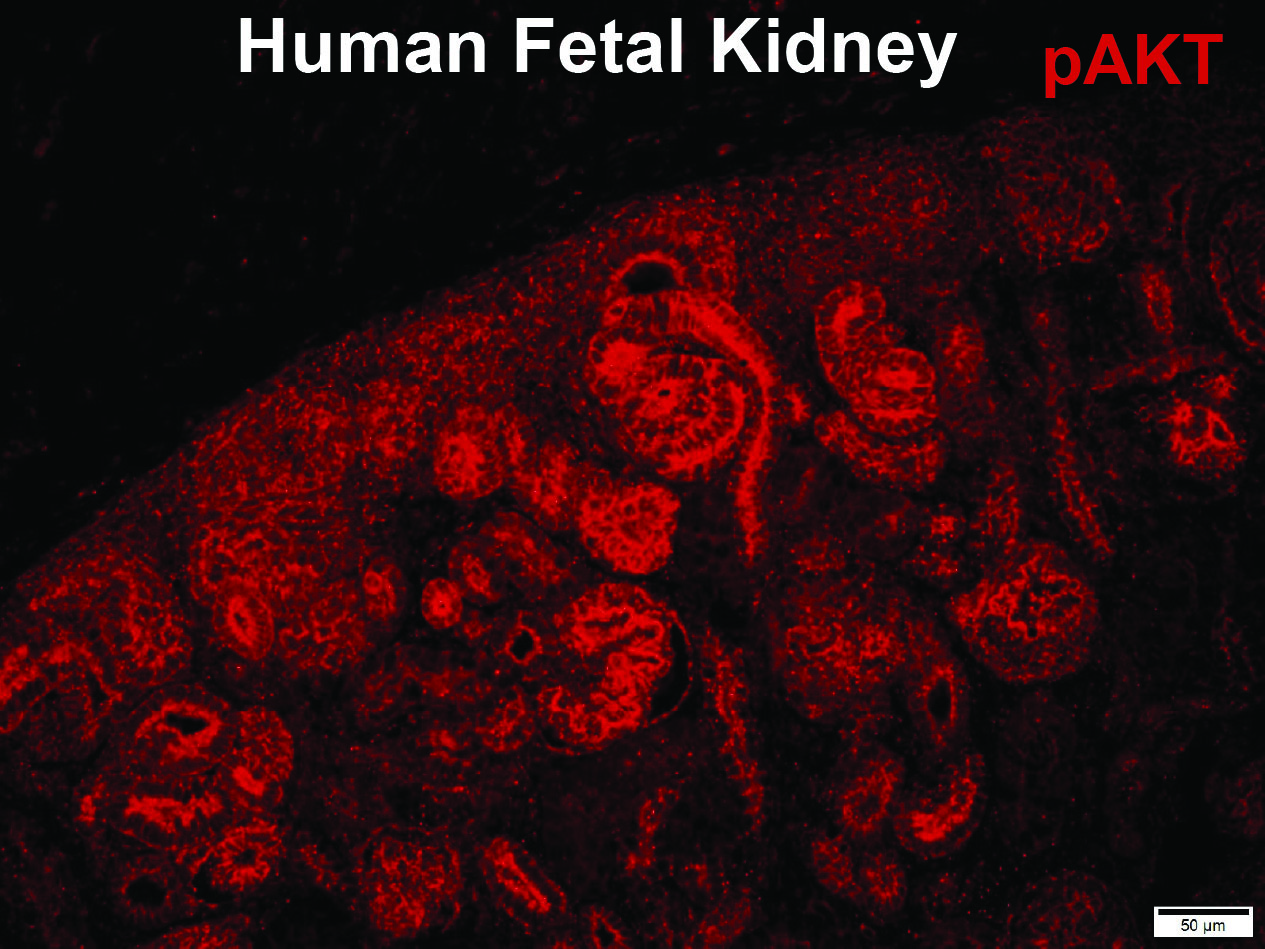

### Supplementary Fig. 16

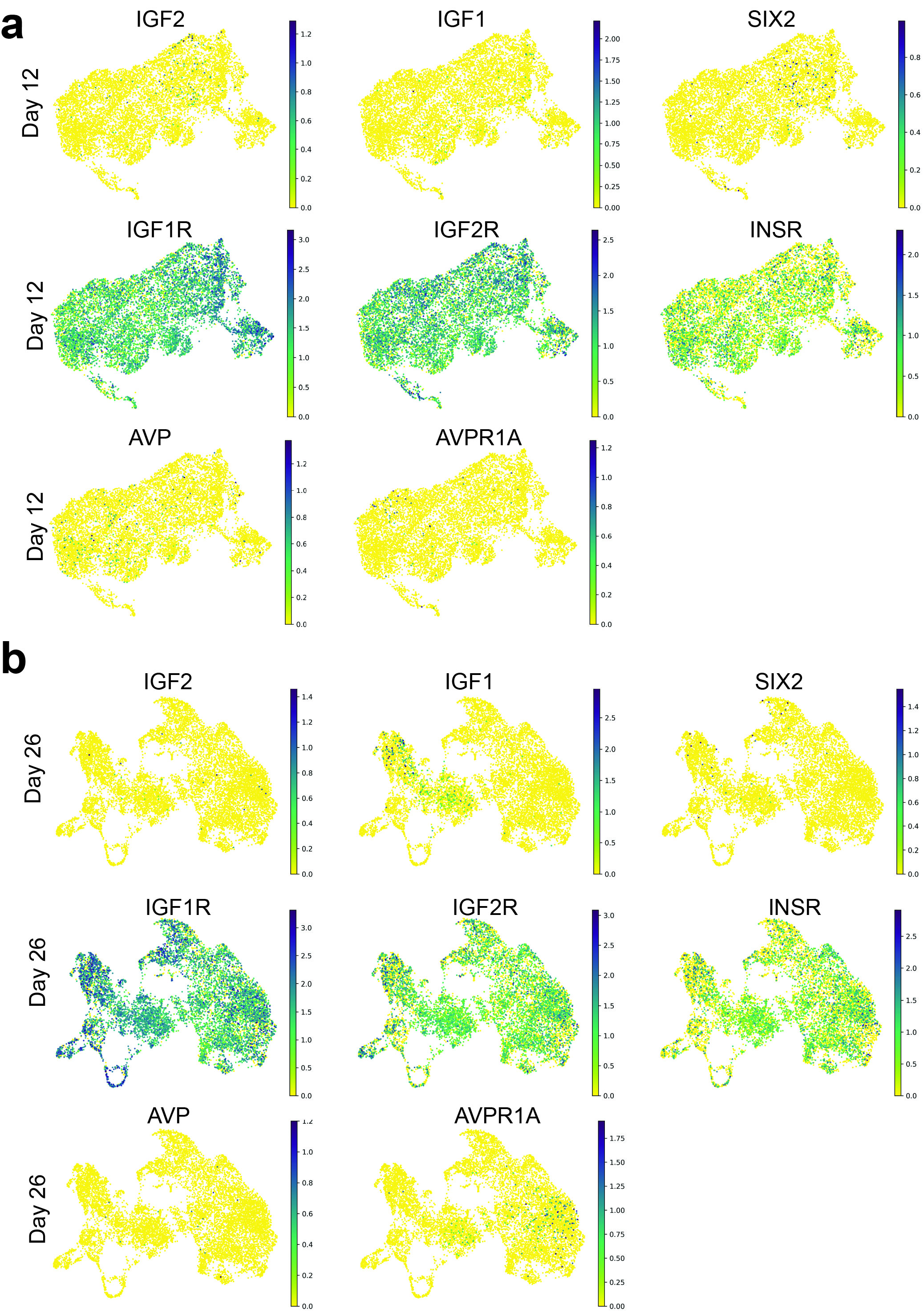

### Supplementary Fig. 17

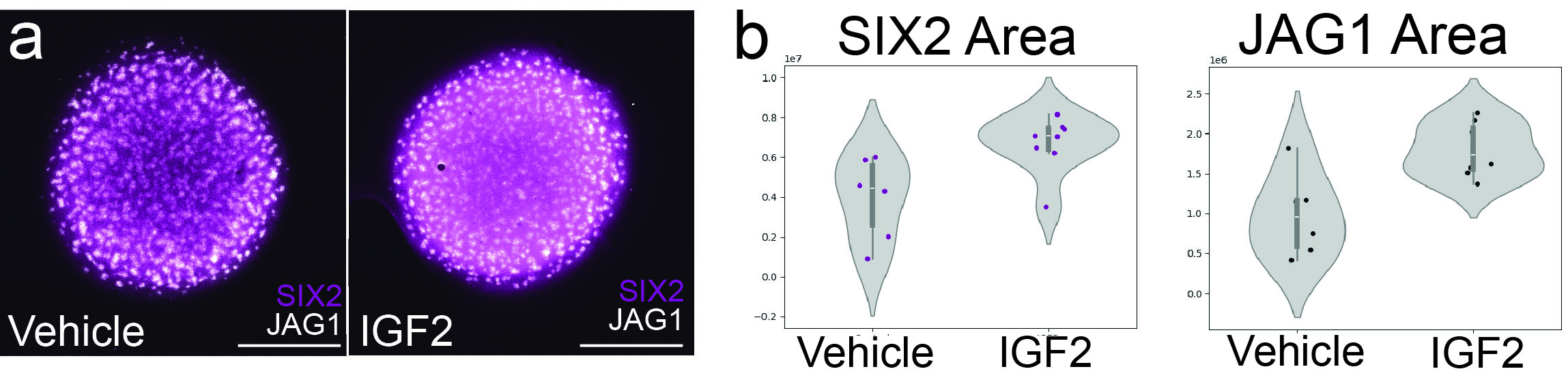

### Supplementary Fig. 18

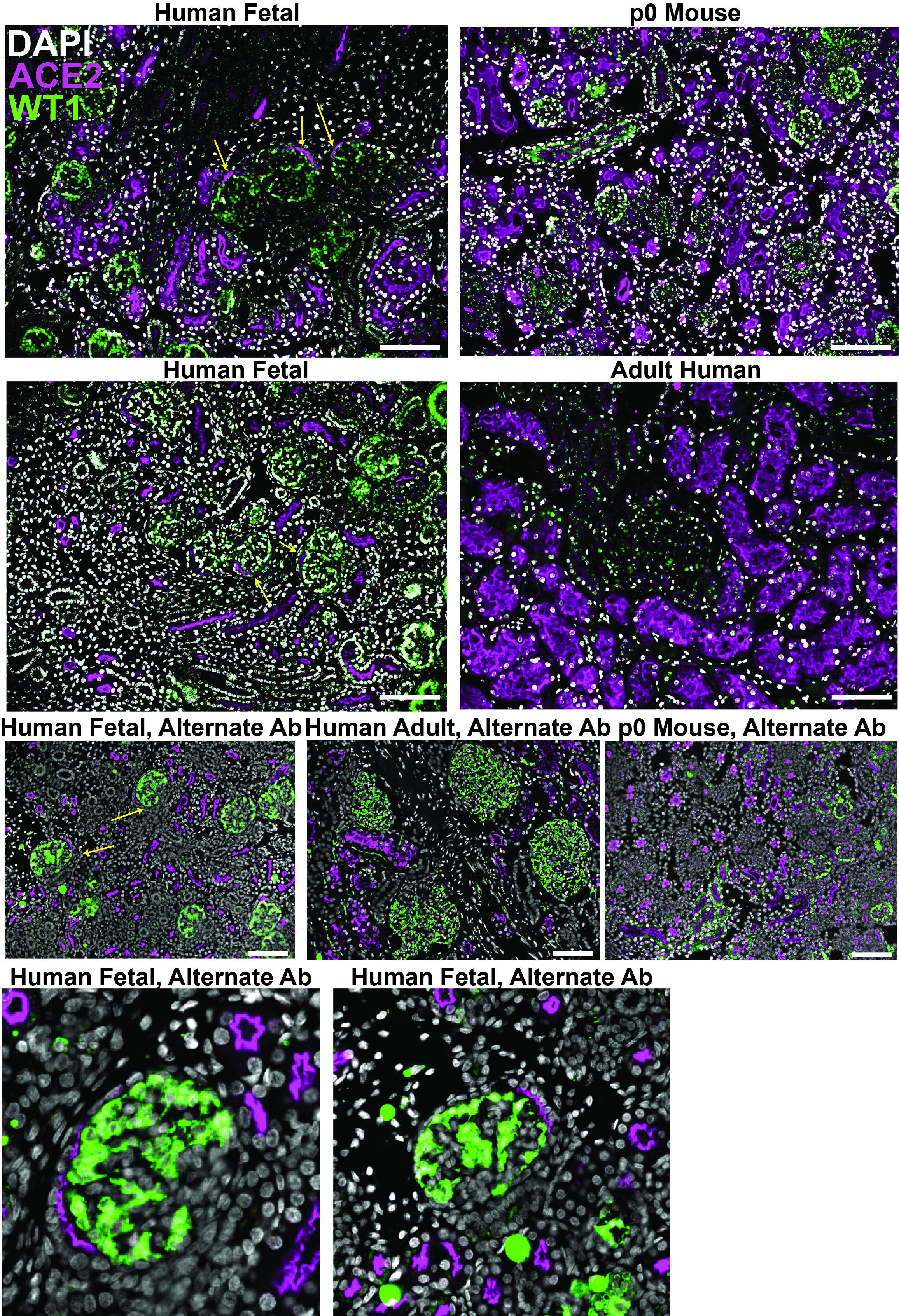

### Supplementary Fig. 19

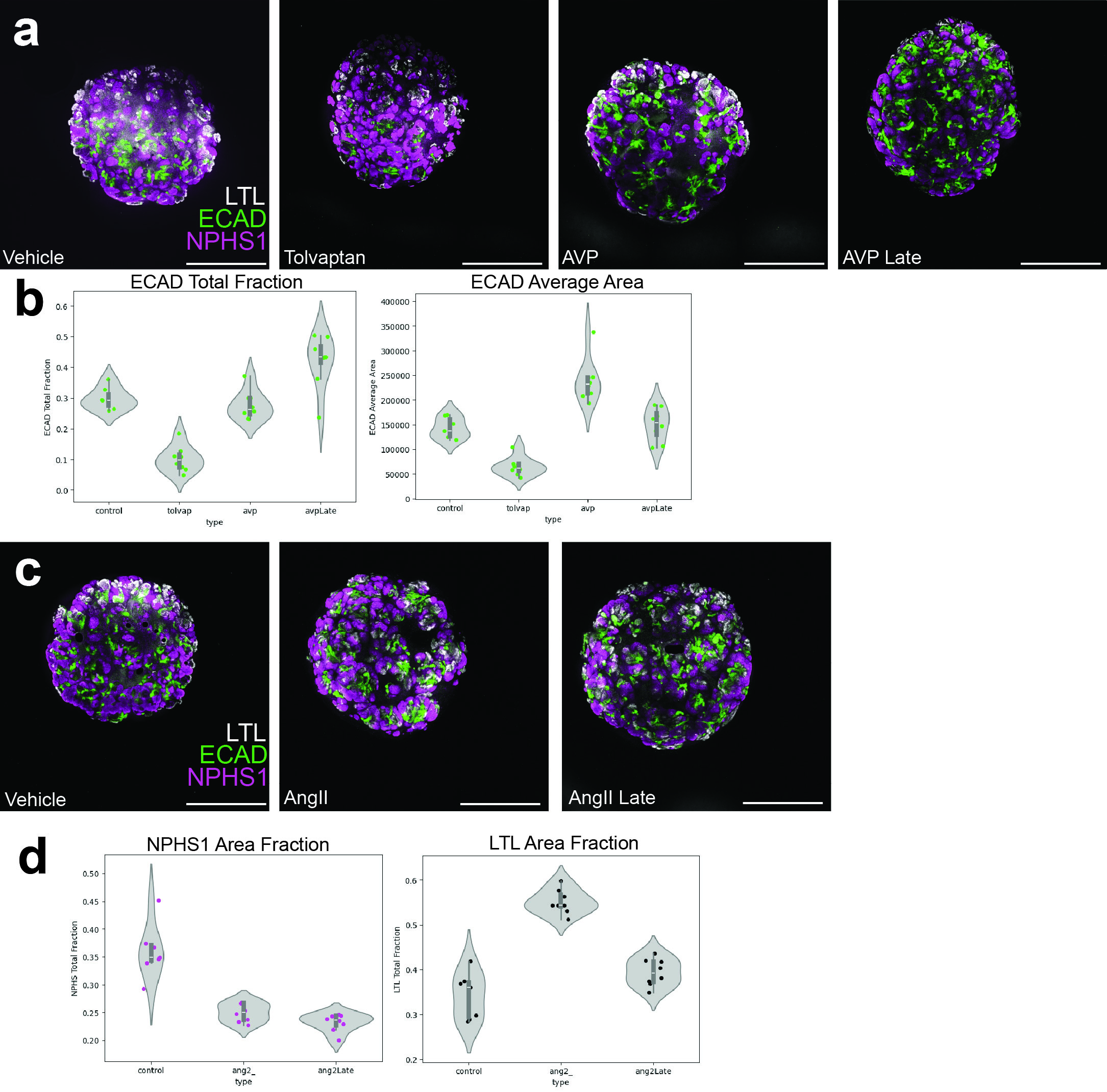

### Supplementary Fig. 20

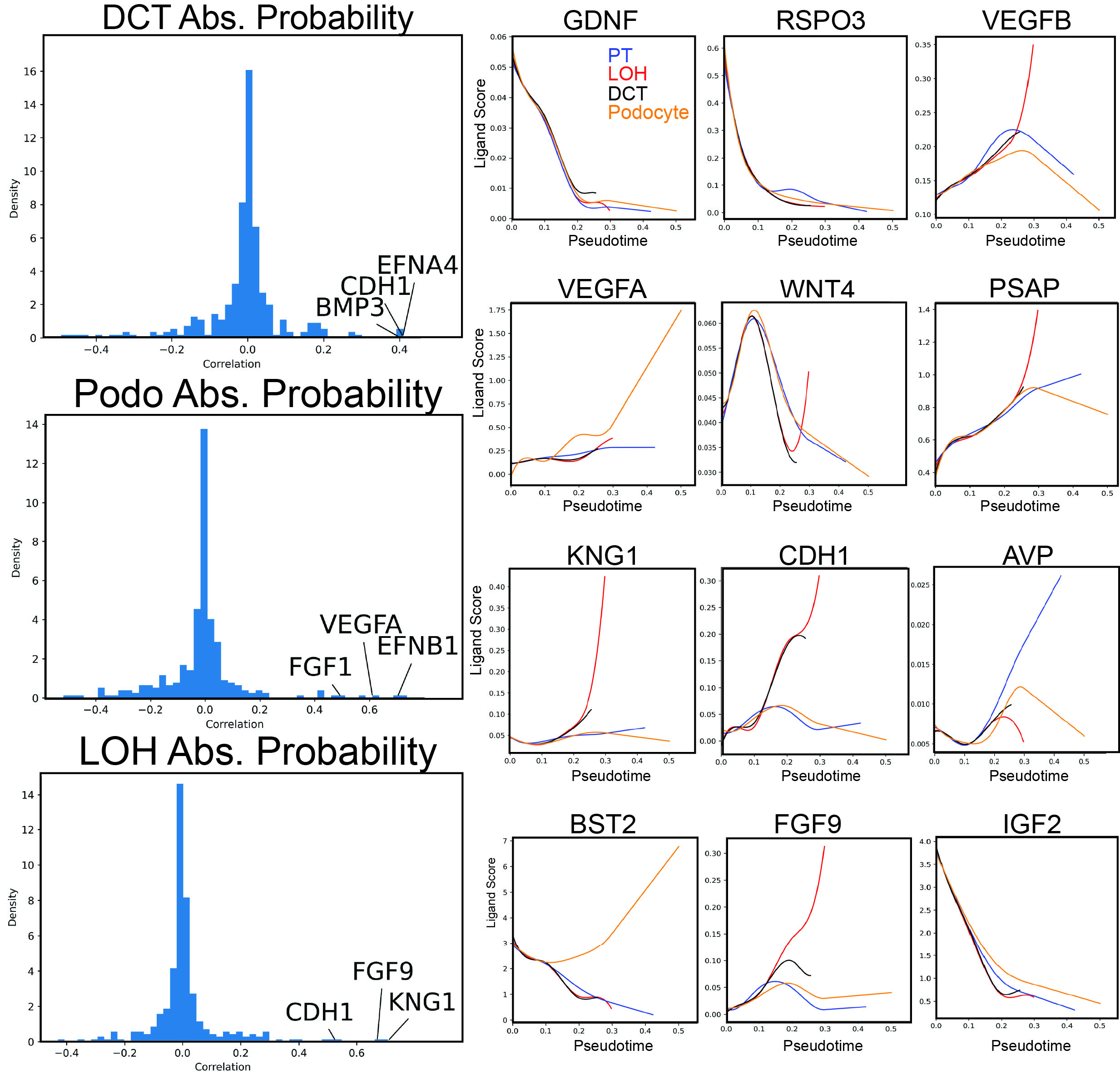

### Supplementary Fig. 22

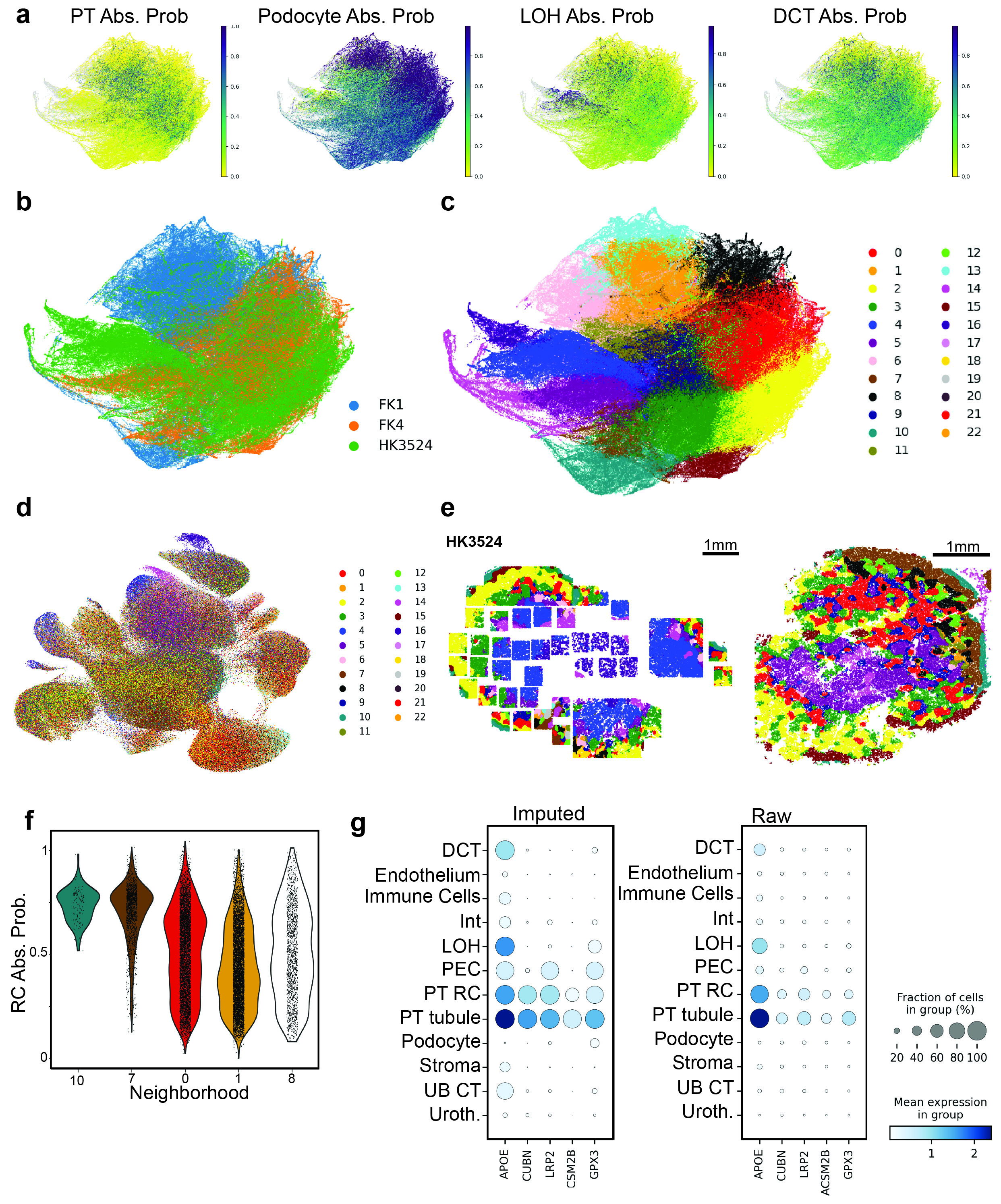
